# Isolation of oxygen-dependent nicotine- and pseudooxynicotine-metabolizing enzymes

**DOI:** 10.64898/2026.08.27.747611

**Authors:** Tejas A. Navaratna, Javeria Akram, Thea D. Pazdernik, Amudha Ramachandran, Pamela Schultz, Mark Dulchavsky, Xavier Choussat, Claire Oczon, Aashnaa Singh, Nikhil Myers, Aaron Robida, Ashootosh Tripathi, Frederick Stull, James C. A. Bardwell

## Abstract

NicA2 is a flavin-bound amine dehydrogenase from *Pseudomonas putida* S16 that converts nicotine to the pharmacologically inactive *N-*methylmyosmine. In animal models of nicotine addiction, injection of NicA2 can decrease nicotine-seeking behavior 10-fold. Accordingly, NicA2-related enzymes have been investigated as smoking-cessation therapeutics. However, efficient catalysis by NicA2 in *Pseudomonas putida* relies on electron transfer to CycN, a cytochrome c, and not directly to O_2_. Impractically high amounts of NicA2 are thus necessary to achieve a pharmacological effect in the absence of CycN. Directed evolution has improved NicA2’s ambient-O_2_ value of k_cat_ from 0.007 s^-1^ to 1 s^-1^, but further improvements have been challenging. Here, we identify a strain of *Peribacillus frigoritolerans* NIC8 which encodes two flavin amine oxidoreductases, Ncox and Pnox. In the presence of oxygen, Ncox and Pnox act on nicotine and pseudooxynicotine respectively with apparent k_cat_ values of 7.7 s^-1^ and 3.9 s^-1^. Transient kinetics establishes bimolecular rate constants of 51100 M^-1^s^-1^ and 81000 M^-1^s^-1^ for the half-reactions between Ncox and O_2_ and between Pnox and O_2_ respectively, consistent with Ncox and Pnox being bona-fide oxidases. Transcriptomics shows enhanced expression of Ncox and Pnox under nicotine-dependent growth as well as supporting the identification of downstream enzymes. Phylogenetic analysis suggests that Ncox and Pnox arose out of repurposing of homologous enzymes found in *Bacillus* species. The enzymes we describe may be useful for the development of nicotine addiction therapeutics and for bioconversion of nicotine in waste streams.

## Introduction

Nicotine-degrading organisms have been isolated from a wide variety of habitats, ranging from the tobacco field rhizosphere^1^, wastewater^2,3^, and tobacco waste extracts^4^. Nicotine is highly toxic, so these strains are of interest due to their bioremediation and bioprocessing potential^5–10^. Nicotine is also highly addictive, and the tobacco use that is driven by nicotine dependence is currently responsible for approximately one in nine deaths worldwide^11^. Consequently, nicotine-degrading organisms are also of great interest as a potential source of enzymes for therapeutic applications. NicA2, a flavoenzyme from *Pseudomonas putida* S16^12^, has been studied in rat models of smoking addiction. Injection of NicA2 into nicotine addicted rats is capable of reducing nicotine-seeking behavior 10-fold^13^, comparing very favorably to the widely utilized nicotine-addiction treatment varenicline (until 2021 marketed as Chantix^®^) which only increases the chance of a successful quit from ∼5% to ∼14%^14^. Like most members of the flavin amine oxidoreductase superfamily, NicA2 first oxidizes a carbon-nitrogen bond, reducing its bound FAD to FADH_2_^15^. To complete the catalytic cycle, FADH_2_ must then react with an electron acceptor. For NicA2 we have previously shown this to be CycN, a cytochrome c found in the periplasm of *Pseudomonas putida*^16^. This requirement makes NicA2 more a dehydrogenase than a true oxidase^17^. NicA2 performs very poorly as a standalone oxidase, as it reacts very slowly with molecular oxygen, exhibiting an apparent k_cat_ value of ∼0.007 s^-1^ ^18^. Its poor oxygen reactivity has necessitated a dosage of up to 70 mg/kg in a rat nicotine addiction model^13^, a therapeutically unrealistic amount for human treatment due to manufacturing, per dose cost, and potential immunogenicity. Rational mutagenesis and directed evolution approaches have been conducted to improve NicA2’s oxygen reactivity, with the most promising variants having an apparent k_cat_ value of ∼ 1 s^-1^ and a K_M_ between 1 and 10 μM. This allowed for the use of a dose of 1 mg/kg to reduce nicotine to undetectable levels in the rat model^19^, more consistent with doses of enzymes currently used for human therapeutics^20^.

However, further efforts to evolve new variants with better k_cat_ values in NicA2 through directed evolution have proven difficult^19^. As an orthogonal approach, we decided to explore the natural evolution and diversity of nicotine degradation pathways through the isolation of organisms from tobacco-related habitats as a source of enzymes with potentially increased functional activity.

Bacteria degrade nicotine through three pathways - the pyrrolidine pathway, carried out by *Pseudomonas* and related Gammaproteobacteria, the pyridine pathway, found in *Arthrobacter-*related organisms, and a hybrid variant pyridine-pyrrolidine (VPP) pathway present in the *Shinella and Agrobacterium* related genera^21^. The pyrrolidine pathway is of primary interest to nicotine degradation for human health applications, as its nicotine degrading enzyme is a relatively small, ∼50 kDa single chain polypeptide that is simple to purify, in contrast to the nicotine hydroxylases present in the pyridine and variant pyridine/pyrrolidine pathways, which are part of a multisubunit, molybdenum-dependent complex, making them likely unsuitable as human therapeutics.

Here we describe the isolation of a strain of *Peribacillus frigoritolerans* we have named *P. frigoritolerans* NIC8, for its ability to grow on nicotine as a sole carbon source. We present detailed genomics and transcriptomics pathway elucidation and mechanistic flavoenzyme characterization. This organism contains a bona fide flavin monoamine oxidase capable of oxidizing nicotine in the presence of oxygen with an apparent k_cat_ of 7.7 s^-1^ and K_M_ of 13.1 μM.

## Results

### Peribacillus frigoritolerans NIC8 can grow using nicotine as the sole carbon source

We isolated a bacterial strain capable of robust growth on M9 agar containing nicotine as its sole carbon source from an unsmoked Marlboro Light cigarette (Figure 1a). The strain failed to grow on M9 agar lacking nicotine. Sequencing of the partial 16S RNA amplicon showed 100% identity to *Peribacillus frigoritolerans* (Supporting Table S1) and FastANI^22^ showed 95.4% average whole-genome nucleotide identity to *P. frigoritolerans* DSM 8801 and alignment fraction of 0.87. The digital DNA-DNA hybridization GGDC formula 2 value of our strain compared with *P. frigoritolerans* DSM 8801 was 63.4%. We therefore have named this newly isolated bacterium *P. frigoritolerans* NIC8, for its ability to grow on nicotine as a sole carbon source. Several other *Peribacillus* strains (YIM B13477, YIM B13472, YIM B13540, YIM B13481, YIM B13482) have recently been reported to grow on nicotine by utilizing the VPP pathway^23^. However, tblastn searches of the genomes of YIM B13472 and YIM B13477 against VPP pathway enzymes *nctB* from *Shinella sp.* HZN7, and *ald, ndhL, ndhS, hsh, and pno* from *Agrobacterium tumefaciens S33* revealed the presence of only fairly distantly related homologs of 22-39% amino acid identity to *P. frigoritolerans* NIC8. Importantly these homologs are also present in *P. frigoritolerans* DSM 8801 and *P. simplex* DSM 1321. *P. frigoritolerans* DSM 8801 and *P. simplex* DSM 1323 failed to grow on nicotine as a sole carbon source (Figure 1b), indicating that growth using nicotine is not a general property of *Peribacillus*.

**Figure 1.**
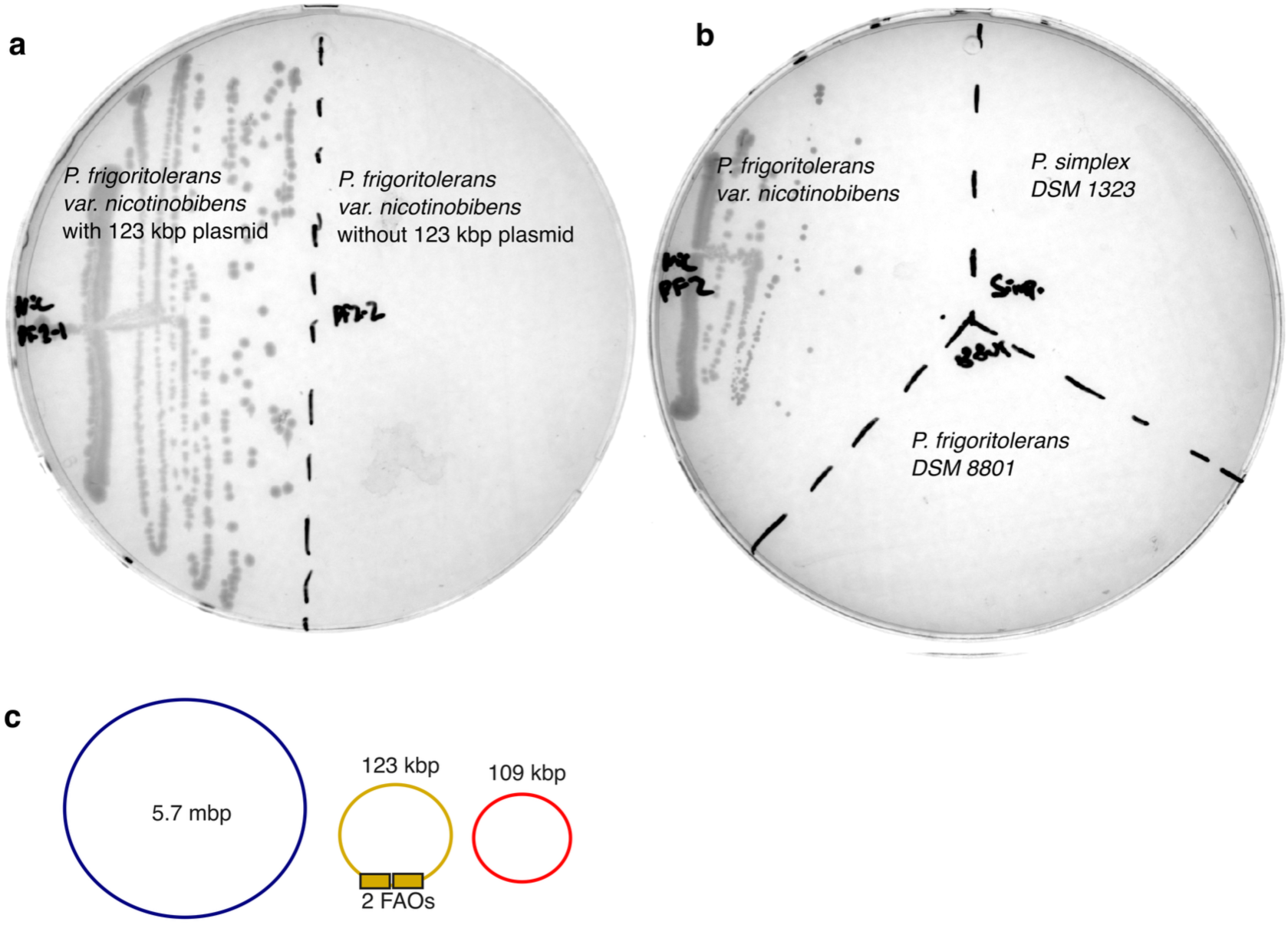
**– a,** Growth of P. frigoritolerans NIC8 as variants having the 123 kbp plasmid pPF-01 present or absent on M9 agarose containing nicotine as the sole carbon source and trace LB medium as a nutrient supplement (0.1% v/v). Plasmid pPF-01 is necessary for robust growth on nicotine. **b,** Growth of Peribacillus strains frigoritolerans NIC8, simplex DSM 1323, and frigoritolerans DSM 8801 The isolate NIC8 is the only one of these three strains capable of using nicotine as the sole carbon source. Plates were imaged after 2 days of growth at 30 °C. **c,** Genome architecture of Peribacillus frigoritolerans NIC8 shows a single 5.7 mbp chromosome and two plasmids of 123 kbp and 109 kbp. The flavin amine oxidoreductases involved in the first two steps of nicotine metabolism are labeled.

### Peribacillus frigoritolerans NIC8 contains a 123-kbp plasmid encoding with two adjacent flavin amine oxidoreductases

Genomic sequencing of *P. frigoritolerans* NIC8 revealed that there were plasmids of 123 kilobases (Kbp) and 109 Kbp in addition to a circular 5.7 Mbp genome (Figure 1c, Supporting Figure S1, Supporting Table S2). The 123 Kbp plasmid pPF-01 was easily cured by passaging of *P. frigoritolerans* NIC8 in LB liquid media overnight, as determined by whole genome sequencing. The resulting plasmid pPF-01-cured strains were unable to grow on nicotine as the sole carbon source (Figure 1a), showing that this plasmid is necessary for nicotine metabolism. The genome of *P. frigoritolerans* NIC8 also contained homologs distant to VPP genes, indicating that these genes do not by themselves support growth on nicotine, consistent with the inability of *Peribacillus* type strains to grow on nicotine plates. Some other nicotine degrading bacteria contain a plasmid responsible for nicotine metabolism including *Shinella sp.* HZN7 and *Paenarthrobacter nicotinovorans*^24,25^; the well-studied nicotine metabolizing bacterium *Pseudomonas putida* S16 contains nicotine degrading genes on a chromosomal island^26^.

Tblastn homology searches using the *P. putida* S16 NicA2 protein as the query sequence against the complete genomic sequence of *P. frigoritolerans* NIC8 showed a pair of open reading frames present on the 123 kbp plasmid pPF-01 with 36.4% and 34.3% identity (Figure 2a) to NicA2 that we termed *ncox* and *pnox*. InterPro searches (ebi.ac.uk/interpro/) confirmed their membership in the flavin amine oxidoreductase protein family (IPR001613) and putative oxidoreductase activity function (GO:0016491), and phylogenetic analysis showed clustering with flavin amine oxidoreductases from Gram-positive bacteria (Figure 2b).

**Figure 2.**
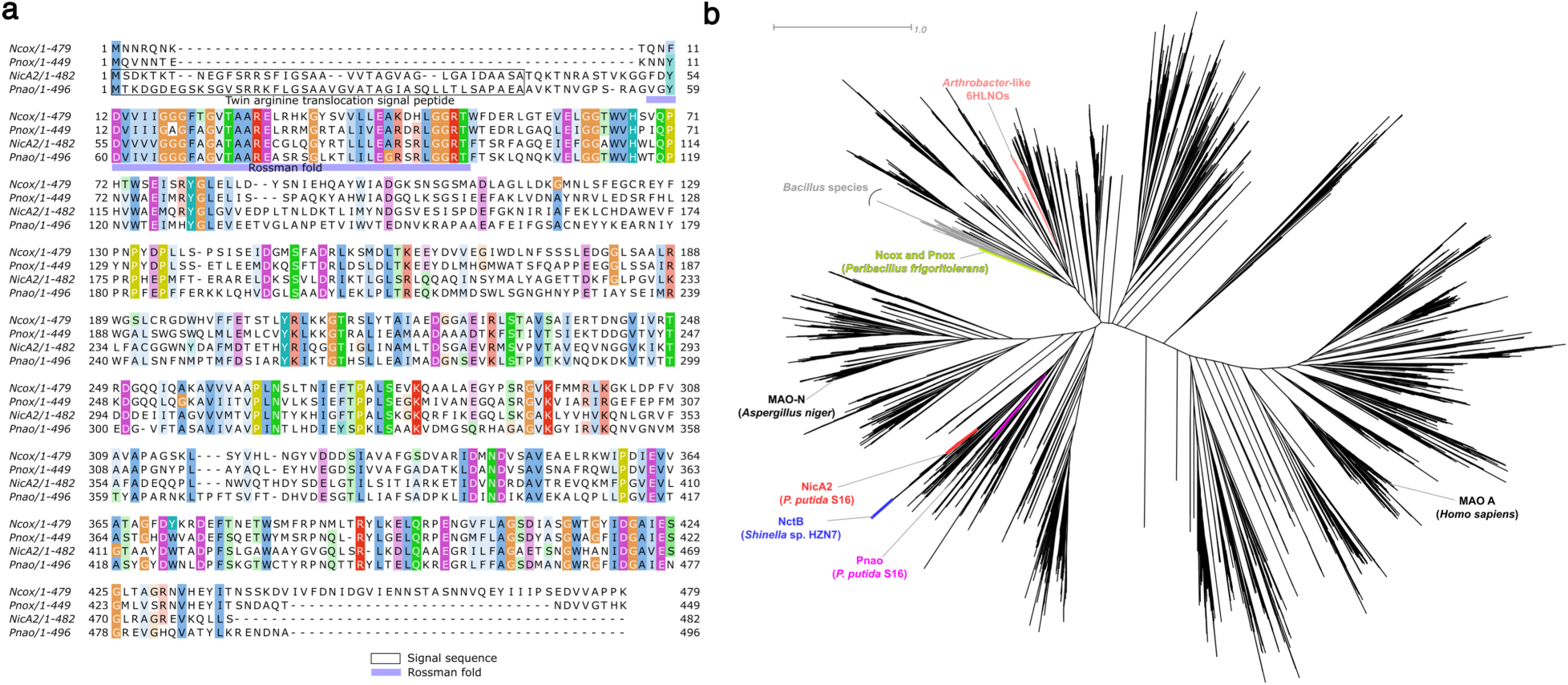
**– a**, Multiple sequence alignment of Ncox and Pnox with Pseudomonas putida S16 NicA2 and Pnao. N-terminal signal sequences were identified with SignalP 6.051. Ncox and Pnox lack an N-terminal twin arginine translocation signal sequence. Colors represent conservation at 25% threshold and amino acid type by the Clustal X color scheme. The highly conserved Rossman fold is apparent from residues 9 – 50 in Ncox (light blue bar). Pairwise identities are as follows: Ncox-Pnox: 54.7%; Ncox-NicA2: 35.2%; Ncox-Pnao: 37.3%; Pnox-NicA2: 33.4%; Pnox - Pnao: 41.0%; NicA2-Pnao: 39.0%. **b**, unrooted cladogram of Ncox and Pnox relative to Pseudomonas putida S16 Pnao and NicA2, Shinella sp. HZN7 NctB, Paenarthrobacter 6HLNOs, and monoamine oxidases (MAOs) from Aspergillus niger and Homo sapiens.

We found that plasmid pPF-01 also contained close homologs to other genes implicated in the pyrrolidine pathway of nicotine metabolism (Figure 3, Supporting Table S3). An aldehyde dehydrogenase gene is located immediately downstream of *pnox.* In the pyrrolidine pathway organism *P. putida* S16, the similarly positioned gene *sapd* encodes an aldehyde dehydrogenase and catalyzes the conversion of 3-succinoylsemialdehyde pyridine to 3-succinoylpyridine^27^ (Supporting Figure S2). The plasmid pPF-01 also contains other pyrrolidine pathway gene homologs ∼20 Kbp away (Figure 4a).

**Figure 3.**
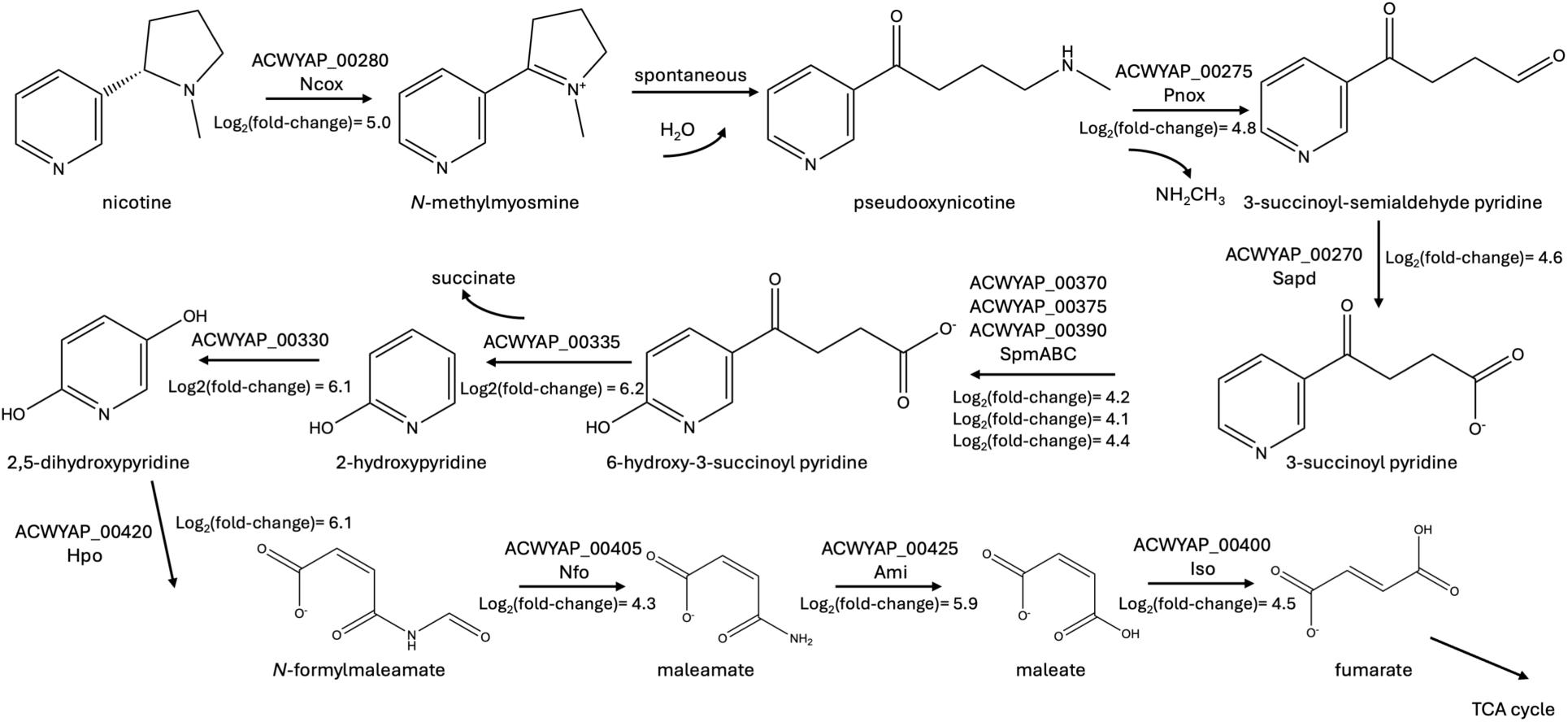
– Proposed pyrrolidine pathway of nicotine metabolism in P. frigoritolerans NIC8. Nicotine is converted to N-methylmyosmine by Ncox, and the spontaneously formed pseudooxynicotine is oxidized by Pnox. Percent identity values to known enzymes are presented in Supporting Table S3.

**Figure 4.**
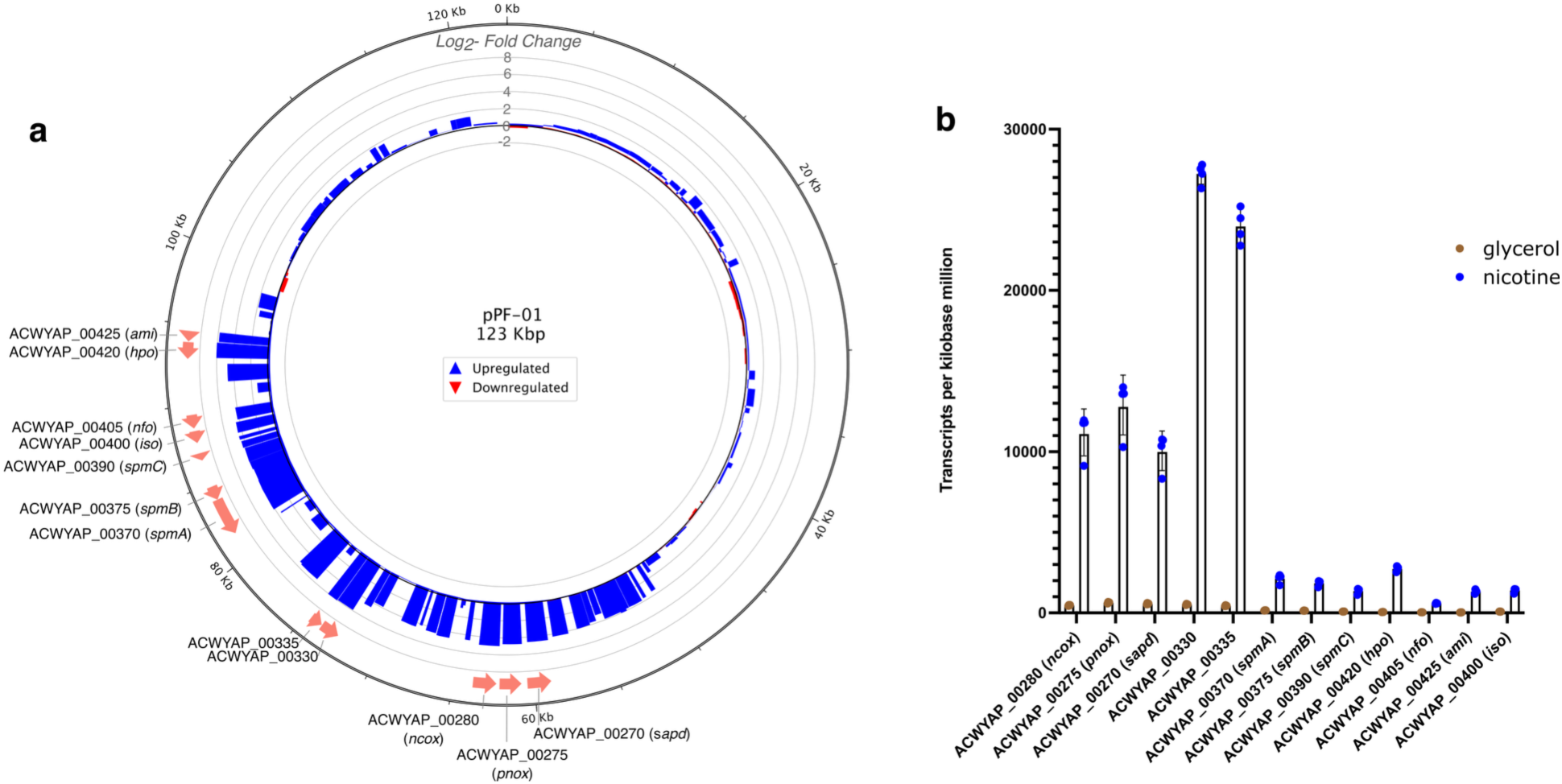
**– a,** Plasmid pPF-01of P. frigoritolerans NIC8 contains nicotine-degrading pathway genes organized in distinct operons. **b**, RNA sequencing analysis of ncox, pnox, and other putative nicotine pathway genes shows upregulation when grown in nicotine as the sole carbon source relative to growth in glycerol.

### Transcriptomics shows nicotine-dependent upregulation of flavin amine oxidoreductases and pathway homologs

*P. frigoritolerans* NIC8 was grown in minimal media containing either glycerol or nicotine as sole carbon source and RNA sequencing was conducted. Differential expression analysis showed that the two flavin amine oxidoreductase homologs *ncox* and *pnox* were highly upregulated during growth on nicotine, with log_2_-fold change values of 5.0 and 4.8 respectively (Figure 3, Figure 4b, Supporting Table S4). The gene *sapd* that lies immediately downstream of *pnox* is also highly upregulated under nicotine growth (log_2_-fold change of 4.6) as are homologs of other pyrrolidine nicotine pathway genes (Figure 3, Supporting Table S4). Genes involved in glycerol metabolism were upregulated in glycerol growth, as expected (Supporting Table S5). Tables of the twenty most upregulated and downregulated genes are presented in Supporting Tables 4 and 5. RNA sequencing coverage mapping of putative nicotine pathway genes shows that the genes are functionally organized in four operons (Supporting Figure S3).

### Kinetic analysis demonstrates rapid O_2_-dependent reactivity and moderate specificity

Zero hits were obtained with a tblastn search using the *P. putida* S16 *cycN* gene^16^ with an E-value of <0.01; a more general HMMER search with PF00034 as the query yielded four hits with an E-value threshold of <0.001. These cytochrome c homologs were all on the chromosome, not the nicotine metabolism plasmid pPF-01, and had log2(fold-change) values of 0.19, 0.47, - 1.5, and 0.16, making them unlikely to be involved in the first step of nicotine conversion compared with log2(fold-change) values in assigned pathway genes in Figure 3 that range from 4.1 to 6.2. Therefore, unlike other bacteria using the pyrrolidine pathway of nicotine catabolism^28^, *P. frigoritoleran*s NIC8 does not encode a recognizable CycN homolog (Supporting Figure S2), suggesting that if a variant of the nicotine pyrrolidine pathway is active in this organism, it has evolved to not utilize this type of electron acceptor. To characterize the enzymatic activity of Ncox and Pnox, these proteins were expressed in *E. coli* and purified (Supporting Figure S4). Solutions of both enzymes were yellow, suggesting the presence of a bound flavin. Trichloroacetic acid precipitation yielded a white precipitate and yellow supernatant, indicating that the flavin is released upon denaturation and that the holoenzyme contains non-covalently bound flavin. Addition of excess nicotine to Ncox under ambient conditions showed rapid change from yellow to clear, indicative of flavin reduction (Figure 5a). No color change of Ncox was observed when excess pseudooxynicotine (the spontaneous hydrolysis product of *N-*methylmyosmine, Figure 3) was added, though a subtle decrease in flavin absorbance may represent protein binding to pseudooxynicotine (Figure 5a), similar to what has previously been reported for NicA2^16^. Conversely, addition of excess pseudooxynicotine to Pnox showed rapid change from yellow to clear, with no absorption change occurring upon excess nicotine addition (Figure 5b). Michaelis-Menten analysis of the reaction of Ncox with nicotine yielded a k_cat_ value of 7.7 s^-1^ and a K_M_ value of 13.1 μM (Figure 5c). Analysis of the reaction of Pnox with pseudooxynicotine found a k_cat_ value of 3.9 s^-1^ and a K_m_ value of 510 μM (Figure 5d). Using the product of Ncox and nicotine instead of commercial pseudooxynicotine yielded similar kinetic constants (Supporting Figure S5). Note that these are apparent values since the velocity measurements were conducted at single oxygen concentration (ambient). This k_cat_ value is ∼1000-fold higher than that observed for NicA2 with oxygen^18^, although the apparent K_M_ of Ncox is ∼100 fold higher than the apparent K_M_ of NicA2 (13.1 μM vs. 0.11 μM).

**Figure 5.**
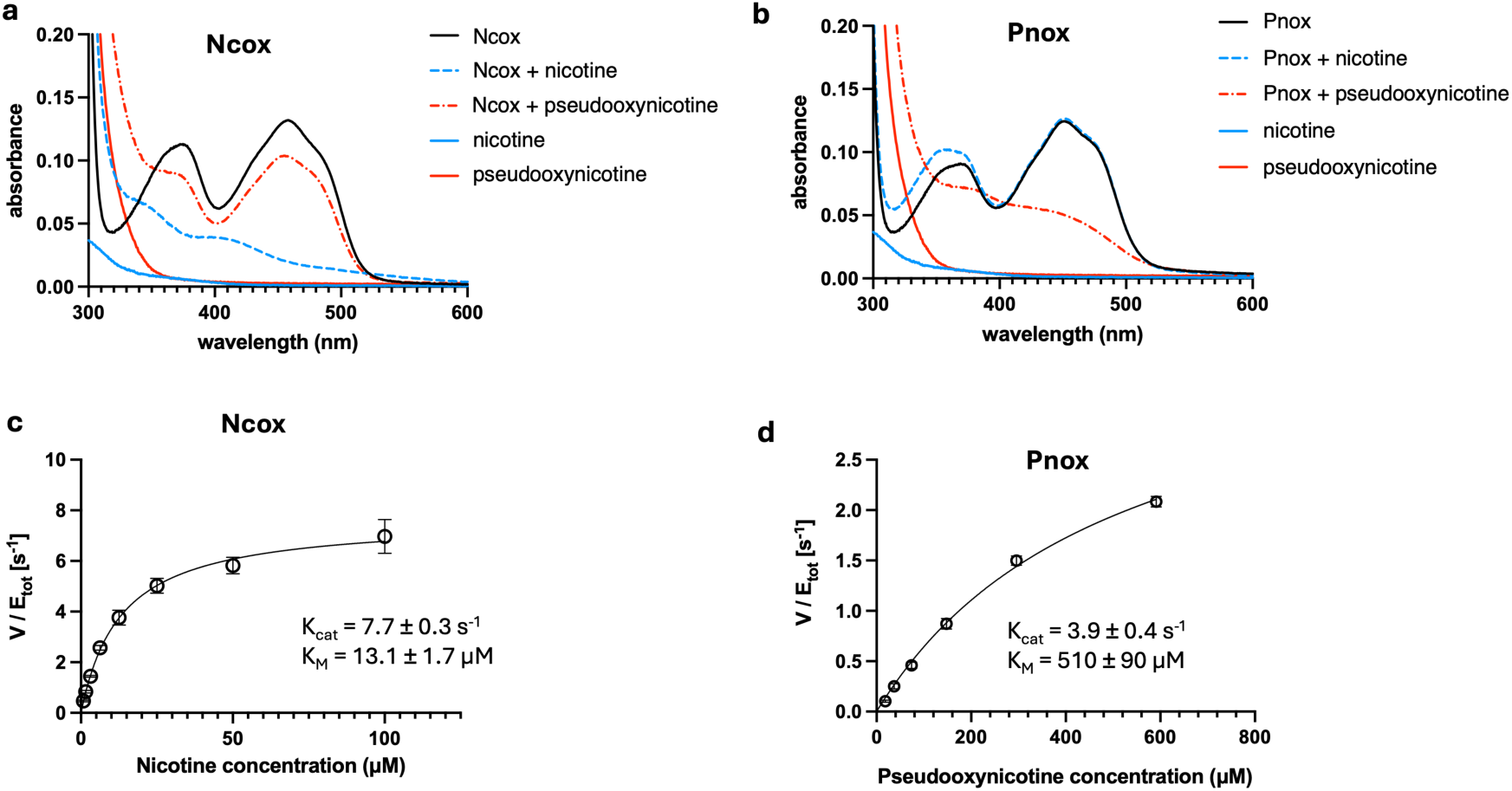
**– a,** UV-vis absorption spectra of Ncox with no substrate (black), excess nicotine (dashed blue), and pseudooxynicotine (dashed red). Addition of nicotine but not pseudooxynicotine results in flavin reduction, which is evident by disappearance of the characteristic 450 nm absorption peak. **b,** UV-vis absorption spectra of Pnox with no substrate (black), excess nicotine (dashed blue), and pseudooxynicotine (dashed red). Addition of pseudooxynicotine but not nicotine results in flavin reduction. **c,** Michaelis-Menten kinetics analysis of Ncox with nicotine as the substrate. The best-fit model parameters are a k_cat_ of 7.7 s^-1^ and a K_M_ of 13.1 μM. **d**, Michaelis-Menten kinetics analysis for Pnox and pseudooxynicotine. Model fitting yielded a k_cat_ value of 3.9 +/- 0.4 s^-1^ and a K_M_ value of 510 +/- 110 μM.

We performed stopped-flow kinetic analysis of flavin absorbance to determine rate constants for each of the half reactions: reduction of flavin by the substrate nicotine or pseudooxynicotine for Ncox and Pnox respectively, and oxidation of the substrate-reduced flavin by molecular oxygen. Ncox’s reduction by nicotine under anaerobic conditions can be described by a triphasic process, in which the fastest phase (k_obs,1_) contributed between 30 – 42% of the overall reaction amplitude (Figure 6a) and the second phase (k_obs,2_) contributed most of the rest of the reaction amplitude. k_obs,1_ showed concentration dependence, with an apparent K_d_ of 53 μM and a reduction rate constant (k_red,1_) of 240 s^-1^ while the second phase had an apparent K_d_ of 18 μM and a rate k_red,2_ of 18 s^-1^ (Figure 6c). A slow third phase was also shown with a rate k_red,3_ of 1.6 s^-1^, contributing 11% of the overall signal change observed.

**Figure 6.**
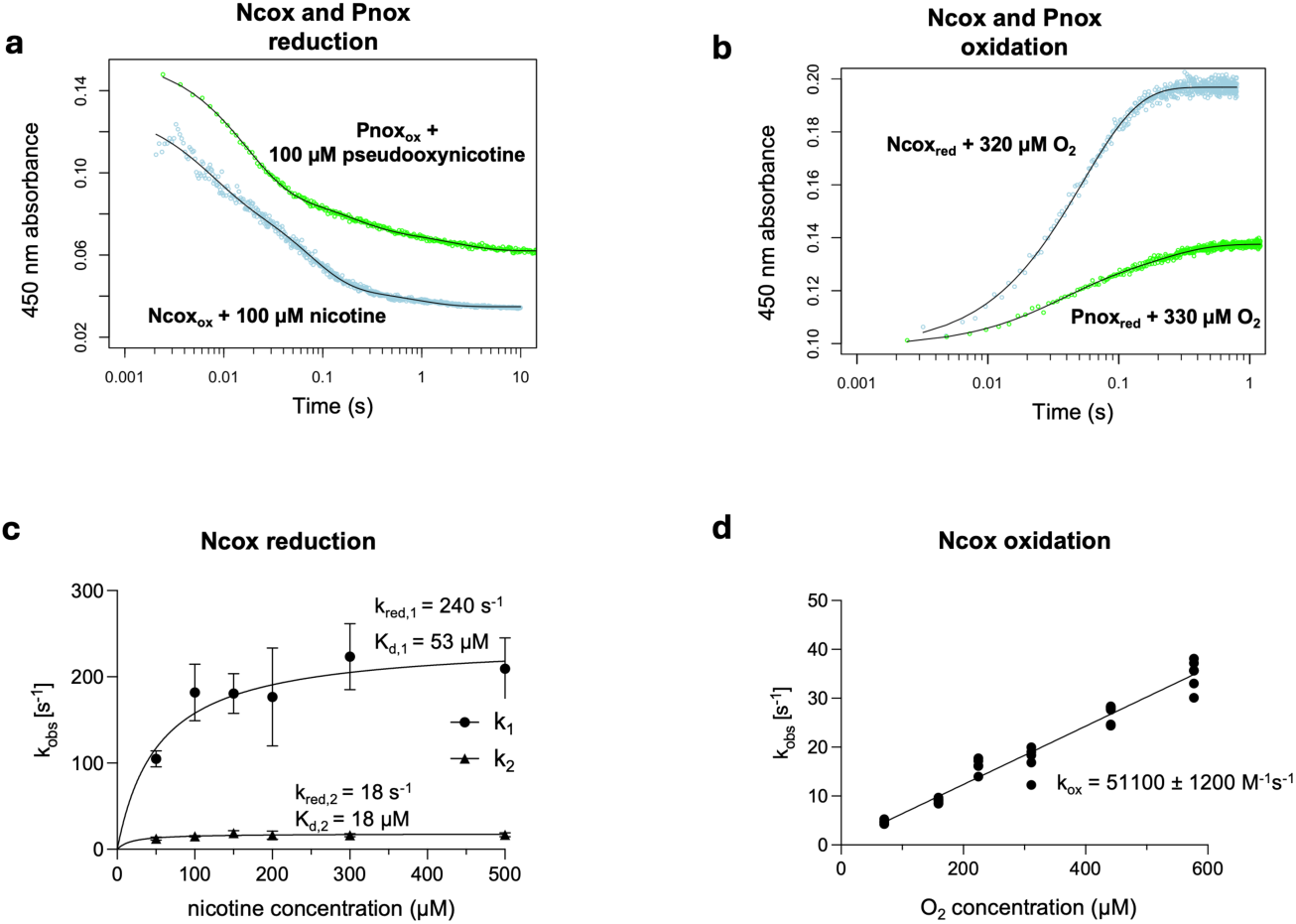
– Half-reaction rate determination. **a,** time curves of Ncox and Pnox reduction by their substrates, nicotine and pseudooxynicotine respectively. **b,** time curves of Ncox and Pnox oxidation by molecular oxygen. **c,** reduction half-reaction rate constants show hyperbolic dependence on nicotine concentration. **d,** oxidation half reaction shows a linear relationship with oxygen concentration, with a second-order rate constant of 51100 M^-1^s^-1^.

Pnox exhibited triphasic reduction behavior upon anaerobic pseudooxynicotine addition. The first phase had a rate k_red,1_ of 170 s^-1^ with an apparent K_d_ of 170 μM and a contribution of 55% - 65% to the overall half-reaction with the second phase having a rate k_red,2_ of 7.4 s^-1^ (Supporting Figure S6a). A slow third phase was observed with a rate k_red,3_ of 0.7 s^-1^, contributing 14% of the signal change. These kinetic parameters are presented in Table 1.

**Table 1.** – Kinetic Parameters. k_red,1-3_ refer to the triple exponential decay rate constants that were fit to absorbance vs. time graphs. K_d,1_ is a fit dissociation constant that describes the relationship between substrate concentration and first exponential reduction rate, as the reduction rate is assumed to be proportional to bound substrate. k_ox,1_ and k_ox,2_ are single exponential (for Ncox) or biexponential (for Pnox) rate constants for oxidation with molecular oxygen. k_cat_, K_M,_ and k_cat_/K_M_ are fit and computed Michaelis-Menten constants. Comparison values for NicA2 were adapted from (a)^19^ or (b)^18^ and values for Pnao were adapted from (c)^31^ or (d)^32^. Ncox and Pnox data were collected at 20 °C. NicA2 and NicA2 v. 320 data were collected at 22 °C^18,19^. Pnao transient kinetics data were obtained at 4°C^31^, whereas steady-state Michaelis-Menten experiments were carried out at 30 °C^32^.

| Enzyme | $k_{red,1}$<br>(s<sup>-1</sup>) | $k_{red,2}$<br>(s<sup>-1</sup>) | $k_{red,3}$<br>(s<sup>-1</sup>) | $K_{d,1}$<br>(μM) | $k_{ox,1}$<br>(M<sup>-1</sup>s<sup>-1</sup>) | $k_{ox,2}$<br>(M<sup>-1</sup>s<sup>-1</sup>) | $k_{cat}$<br>(s<sup>-1</sup>) | $K_M$<br>(μM) | $k_{cat}/K_M$<br>(M<sup>-1</sup>s<sup>-1</sup>) |
| --- | --- | --- | --- | --- | --- | --- | --- | --- | --- |
| Ncox | 240 | 18 | 1.6 | 53 | 51000 | N/A | 7.7 | 13 | 5.9 x 10<sup>5</sup> |
| Pnox | 170 | 7.4 | 0.7 | 170 | 81000 | 15000 | 3.9 | 510 | 7.6 x 10<sup>3</sup> |
| NicA2 | 800<sup>a</sup> | 120<sup>a</sup> | - | 31<sup>a</sup> | 28<sup>a</sup> | - | 0.0061<sup>b</sup> | 0.11<sup>b</sup> | 5.4 x 10<sup>4</sup><sup>b</sup> |
| NicA2 v.320 | 19<sup>a</sup> | 4.1<sup>a</sup> | - | N.D | 4,400<sup>a</sup> | - | 1.1<sup>a</sup> | 6.3<sup>a</sup> | 1.7 x 10<sup>5</sup><sup>a</sup> |
| Pnao | 74<sup>c</sup> | 6<sup>c</sup> | - | 64<sup>c</sup> | 600<sup>c</sup> | - | 0.79<sup>d</sup> | 73<sup>d</sup> | 1.1 x 10<sup>4</sup><sup>d</sup> |

Both enzymes Ncox and Pnox therefore had two major kinetic phases that corresponded to flavin reduction. This phenomenon has been observed for the homologous enzymes NicA2^16^ and NctB in *Shinella sp.* HZN7 which oxidizes (*S*)-6-hydroxynicotine into 6-hydroxy-*N*-methylmyosmine. In both cases these two phases have been interpreted as indicating differential reactivity by the two subunits of the homodimeric enzyme.

Reaction of both Ncox and Pnox with oxygen was found to be rapid, with a second order rate constant (k_ox,1_) of 51100 +/- 1200 M^-1^ s^-1^ for Ncox and 81000 +/- 20000 M^-1^ s^-1^ for Pnox. Pnox, but not Ncox, was better fit with a biexponential process, indicating two sequential kinetic steps with roughly equal kinetic amplitudes (Figure 6b, Supporting Figure S6b). This may be a consequence of differing oxidation rates for product-bound vs. free enzyme species, as Pnox enzyme was initially reduced with its substrate pseudooxynicotine. Interestingly, k_ox_ for Ncox was unaffected by the presence of 2 mM pseudooxynicotine (Supporting Figure S7), the product of nicotine oxidation. Therefore, Ncox is somewhat different from the k_cat_-enhanced NicA2 variant v.320, which exhibits greatly accelerated reaction with oxygen in the presence of excess pseudooxynicotine^19,29^. These k_ox,1_ values for Ncox and Pnox are similar to those seen for characterized *bona fide* flavin monoamine oxidases such as human spermine oxidase^30^ (40000 M^-1^ s^-1^) and *Shinella sp.* HZN7 NctB^28^ (38,000 M^-1^ s^-1^) but much faster than for NicA2 for which this reaction rate is 28 M^-1^ s^-1^ ^19^. Since the rate at which NicA2 is reduced by CycN is much more rapid at 1 x 10^6^ M^-1^ s^-1^, NicA2 is better characterized as a dehydrogenase than as an oxidase. Additionally, the Amplex® Red assay confirmed near-stoichiometric production of H_2_O_2_ by the Ncox-nicotine reaction and Pnox-pseudooxynicotine reactions (Supporting Figure S8). Half-reaction kinetics data are presented in Table 1.

### Ncox allows robust growth in a Pseudomonas cycN deletion strain

Because these results suggest that Ncox’s natural electron acceptor is molecular oxygen, we probed if Ncox can support oxygen and nicotine-dependent growth of the *P. putida* S16 strain that has had its *cycN* gene deleted. This strain normally exhibits very poor growth on nicotine plates, a phenotype that we have previously used to select for oxygen utilizing variants of NicA2^16,19^. Our most effective selected oxygen utilizing variant of NicA2 (NicA2v320) has a k_cat_ value for nicotine in the presence of oxygen of 1.1 s^-1^. Ncox has a k_cat_ value for the conversion of nicotine in the presence of oxygen of 7.7 s^-1^, 7-fold better. Consistent with these values, Δ*cycN P. putida* S16 can be complemented better by Ncox than it can by NicA2v320 (Figure 7). SDS-PAGE analysis shows that Ncox is expressed to similar levels as NicA2v320 (Supporting Figure S9); therefore, faster growth of the Ncox-expressing strain is likely attributable to the increased catalytic rate of Ncox vs. NicA2v320.

**Figure 7.**
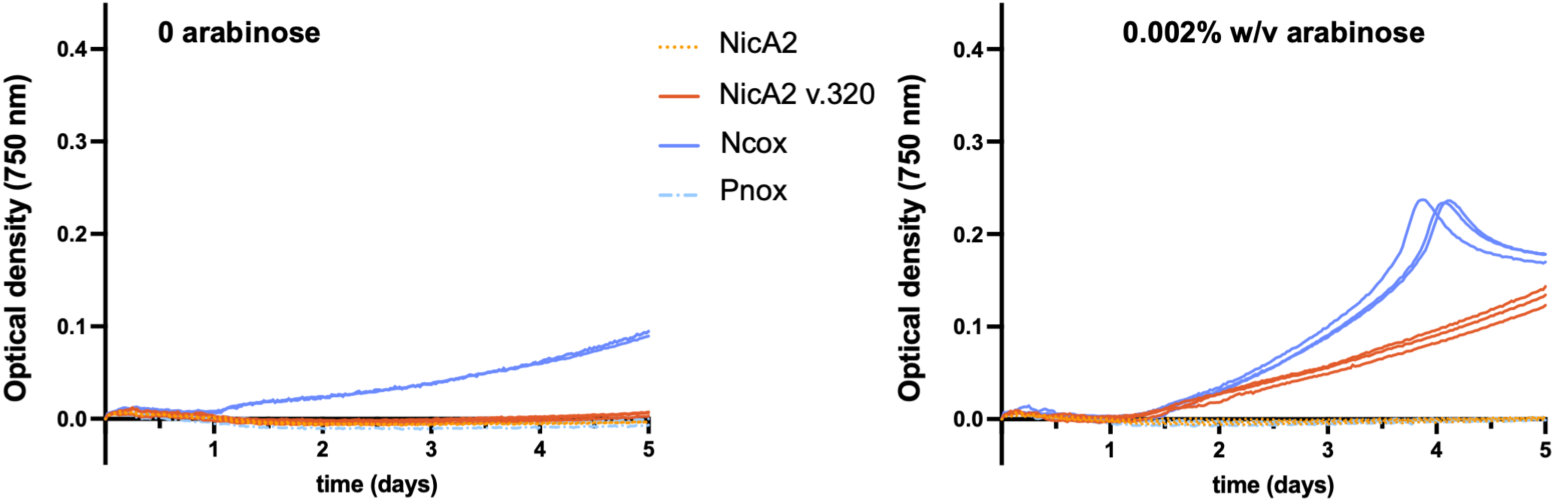
– Growth of P. putida S16 ΔcycN complemented with nicotine degrading enzymes shows that leaky expression alone (0 arabinose) allows for growth with Ncox, but no other enzyme. At an arabinose concentration of 0.002%, Ncox confers a higher growth rate than the fastest growing NicA2 variant v.320. In each plot, all three replicates are shown.

## Discussion

Due to the interest in developing nicotine-degrading enzymes as addiction therapeutics, there is demand for enzymes that are able to rapidly degrade nicotine. One potential therapeutic is NicA2, from *Pseudomonas putida* S16^13,33,34^. However this enzyme has a very low apparent k_cat_ of 0.007 s^-1^ under ambient conditions when using oxygen as an electron acceptor, as it relies on the cytochrome c CycN for activity in *Pseudomonas putida* S16^16^. This is clearly not ideal for use as a stand-alone pharmaceutical enzyme. Therapeutic supplementation with CycN would be challenging as this would likely also require co-supplementation with a cytochrome c oxidase, which are large integral transmembrane protein complexes, in order to have multiple enzyme turnover events. Subsequent efforts to improve NicA2’s reaction rate with oxygen through design and screening improved this k_cat_ to 0.16 s^-1^ ^35^, and our lab’s previous directed evolution experiments resulted in improvements to 1.1 s^-1^ ^19^. Ongoing research with NicA2 and close homologs has failed to improve apparent k_cat_ values beyond this point, so as an alternative approach we looked to natural microbial evolution. By prospecting for organisms that could utilize nicotine as the sole carbon source, we found the distant NicA2 homolog, Ncox, which breaks down nicotine to the pharmacologically inactive compound *N*-methylmyosmine in a directly-oxygen-dependent manner. Ncox has a k_cat_ seven-fold higher than our best laboratory-evolved NicA2 variant and by this criterion, is an attractive candidate as a nicotine degrading therapeutic enzyme.

Reducing power, harnessed from oxidizing various substrates, is generally useful to bacteria. Gram-negative bacteria use a membrane-bound electron transport chain for aerobic respiration and ATP generation. All previously characterized nicotine-specific FAD oxidoreductases are in the bacterial pyrrolidine pathway, that we are aware of, and these enzymes including NicA2, have a cytochrome c gene closely linked^16^. These NicA2-related proteins contain an apparent signal sequence (Figure 2a). This strongly suggests that these NicA2 homologs are localized to the periplasmic space and rely on a cytochrome c homolog as an electron acceptor. In contrast, 6-hydroxy-L-nicotine oxidases (6HLNOs) which act on (*S*)-6-hydroxynicotine in the pyridine pathway use molecular oxygen, lack a periplasmic localization signal, and can be found in gram-positive bacteria including *Arthrobacter*^36^. Unlike periplasmically localized NicA2, Pnao, and homologs in *Pseudomonas*, Ncox and Pnox lack a twin-arginine-translocation signal sequence (Figure 2a), and are likely cytoplasmic in the Gram-positive bacterium *P. frigoritolerans* NIC8.

Recently, we showed that FAD oxidoreductases have made the switch multiple times between the common default of utilizing oxygen and utilizing alternative electron acceptors such as cytochrome c^37^. Through ancestral sequence reconstruction we showed in particular that a common ancestor of *Shinella* sp. HZN7 6HLNO and NicA2 was likely to have used a cytochrome c, and that oxygen utilization only appeared in the *Shinella* 6HLNO branch after the split with NicA2^37^. Moreover, the 6HLNO found in *Shinella sp.* HZN7 is evolutionarily distant from the 6HLNO found in *Paenarthrobacter nicotinovorans*^293029^ (Figure 2b), suggesting that metabolism of (S)-6-hydroxynicotine^21,29^, and by extension, of nicotine can be evolved separately multiple times. The amino acid sequences of the two adjacent flavin amine oxidoreductases in *P. frigoritolerans* NIC8 are more similar to each other (55% identity) than they are to any known nicotine catabolic pathway flavin amine oxidoreductase (max. 41%, Pnox vs. Pnao in *P. putida* S16). This suggests that Ncox and Pnox evolved from a common ancestral flavin amine oxidoreductase, likely within the bacillus clade (Figure 2b), and arose independently of any nicotine-related enzymes previously described. Structurally, there are homologous aromatic cage residues in Ncox as in NicA2: NicA2’s W427 aligns to W381 in Ncox, W108 in NicA2 is W65 in Ncox, whereas W364 in NicA2 is Y319 in Ncox; this aromatic cage is highly conserved in flavin amine oxidoreductases and is important for oxidation of nicotine^18^. However, NicA2’s T381 and N462 are replaced by F335 and Y417 respectively in Ncox (Figure 2a), which may contribute to additional pi-pi stabilization of nicotine or interaction with FAD’s isoalloxazine ring. Understanding this local chemical environment, how product is released, and oxygen accessibility for Ncox and Pnox are areas of future study.

Tying nicotine breakdown to the electron transport chain allows *Pseudomonas* and related organisms to extract considerable reducing power from nicotine and this advantage likely explains the seeming lack of naturally evolved cytochrome c-independent, oxygen dependent NicA2 variants. NicA2, Pnao and their close homologs are periplasmically localized in Gram-negative bacteria, consistent with the interplay of the inner-membrane bound cytochrome and periplasm-localized redox enzyme systems^38^. In contrast, all previously described nicotine-degrading Gram-positive bacteria we are aware of utilize the pyridine pathway to degrade nicotine with a cytoplasmically localized multisubunit nicotine hydroxylase enzyme complex^39^. These complexes are generally difficult to purify and contain molybdenum cofactors making them much less tractable pharmacologically than the homodimeric NicA2 homologs. We present here a report of a Gram-positive bacterium that utilizes the pyrrolidine pathway to degrade nicotine and uses molecular oxygen as the direct oxidant of the FAD cofactor in the first two steps of nicotine catabolism. The enzyme Ncox from *P. frigoritolerans* NIC8, like its homolog NicA2 from *Pseudomonas putida* S16^12^ breaks down nicotine to the pharmacologically inactive compound NMM. Despite Ncox’s larger oxygen-dependent k_cat_ value over NicA2 and fastest directed evolution NicA2 variants (7.7 s^-1^ vs. 1.1 s^-1^)^19^, the k_cat_/K_M_ or catalytic efficiency of all of these enzymes for nicotine degradation are between 10^5^ to 10^6^ M^-1^ s^-1^. Ncox has a k_cat_/K_M_ of 6 x 10^5^ M^-1^ s^-1^. Across the broad range of flavin amine oxidoreductases, many have k_cat_/K_M_ values of maximally 10^6^ M^-1^ s^-1^ ^40,41^. One proposed reason for this is a k_cat_-K_M_ tradeoff. Flavin amine oxidoreductases typically undergo a ping-pong kinetic mechanism, in which the bound flavin is reduced, thereby oxidizing its substrate, and then, a subsequent step occurs in which the flavin is oxidized. With this mechanism, k_ox_ contributes to both the apparent k_cat_ and apparent K_M_ for nicotine such that they should maintain the same proportion (kcat/Km) if k_ox_ changes^42^. This likely contributes to why Ncox has the highest apparent K_M_ for nicotine since it has the highest k_cat_ and k_ox_. If this explanation for the observed consistency in k_cat_/K_M_ is correct, any improvements in k_cat_ through oxygen reactivity via natural or laboratory evolution may be abrogated through an increased K_M_. Indeed, our lab previously reported an experimental plateau of k_cat_ ∼ 1.3 s^-1^ and K_M_ ∼ 1 μM for directed evolution of NicA2^19^. However, this directed evolution plateau may merely be a consequence of the high concentrations of nicotine used in selections, i.e. a lack of evolutionary pressure on K_M_. In smokers, the peak plasma nicotine concentration varies between 10-50 ng/mL^43^ (60 – 300 nM) which is substantially lower than the K_M_ of both NicA2 v.320 and Ncox. The conversion rate is therefore governed by k_cat_/K_M_. Improvements to the K_M_ are as important as improvements in k_cat_ and it remains to be seen if mutations can substantially increase the catalytic efficiency of Ncox.

## Experimental procedures

### Isolation of Peribacillus frigoritolerans NIC8

Samples were collected from different nicotine-rich sources, including tobacco leaves from cigars, commercial cigarettes, and soil from outside smoking areas. Cigars and cigarettes were cut by sterilized scissors to collect the internal tobacco leaves. Initial screening was performed by inoculating samples onto NPD nicotine agar plates (6.0 g/L Na_2_HPO_4_, 3.0 g/L KH_2_PO_4_, 0.5 g/L NaCl, 1.0 g/L NH_4_Cl, 0.2 g/L MgSO_4_ heptahydrate, 0.01 g/L CaCl_2_, 0.05% nicotine v/v, 1 mL/L trace element solution, and 1.5% w/v agar) at 28°C for 14 days. Nicotine degrading microbial colonies were selected based on visible growth difference between NPD nicotine agar plates and NPD agar plates lacking nicotine.

Among resulting isolates, several with well described nicotine degradation pathways were identified (e.g. *Pseudomonas* and *Paenarthrobacter* strains). A previously undescribed additional isolate from a Marlboro Light cigarette stored in a cigarette case exhibited robust growth on an NPD nicotine agar plate. A single colony of this isolate was transferred onto NPD nicotine agar serially 5 times to ensure axenic purity. A 16S amplicon was generated from PCR using whole cells with primers 5’-AGAGTTTGATCMTGGCTCAG and 5’-CGGTTACCTTGTTACGACTT and Sanger sequencing was carried out at Azenta using primer 5’-AGAGTTTGATCMTGGCTCAG. The resulting read was trimmed at both ends by quality, resulting in a 923 bp query (Supporting Table S1) which was analyzed using the NCBI BLAST webserver using the 16S ribosomal RNA sequences database.

### Genome sequencing and PCR

10 mL of liquid M9-traceLB-nicotine (6 g/L Na_2_HPO_4_, 3 g/L KH_2_PO_4_, 1 g/L ammonium chloride, 1 mM MgSO_4_, 0.1 mM CaCl_2_, 1× trace metals (Teknova, T1001), 1 μg/mL thiamine, 0.5 g/L nicotine, and 0.125% v/v LB medium) was inoculated with a single colony from an M9-traceLB-nicotine agarose plate (M9-traceLB-nicotine with 15 g/L agarose). Following overnight growth at 200 rpm at 30 °C, cells were spun down, the pellet was resuspended in 500 μL Zymo DNA/RNA Shield and sent to Plasmidsaurus for standard bacterial DNA extraction and sequencing as follows: approximately 275000 reads were gathered, followed by base-calling with Dorado v4.3 and quality filtering with Filtlong v0.2.1 to remove reads shorter than 300 bp and keeping highest quality 95%. Flye v2.9.1^44^ was used to generate an assembly using parameters optimized for high-quality ONT reads and the assembly was polished with Medaka v1.8.0.

For plasmid curing, a single colony of *P. frigoritolerans* NIC8 was transferred serially 5 times in 5 mL liquid LB media, followed by streaking out on an LB agar plate and colony PCR using primers F-CTCAAGGAGGTTTTTAAATGCAAGTGAACAATACAG and R-TAGAACAATTCGATTATTTATGCGTACCAACAACGTC to confirm absence of *pnox* by lack of the 1380 bp band corresponding to the *pnox* gene. *pnox^-^* colonies were inoculated into 3 mL LB liquid media and grown overnight at 200 rpm at 30 °C. Upon validating lack of *pnox* by PCR, pellets were resuspended in 500 μL Zymo DNA/RNA Shield and sent for standard bacterial DNA extraction and whole genome sequencing through Plasmidsaurus to confirm curing.

### Homology searches, sequence comparisons, and phylogenetics

For tblastn searches for homologs, *P. frigoritolerans* DSM 8801 and *P. simplex* DSM 1321 genomes were downloaded from GenBank via accessions GCF_024169475.1 and GCF_002243645.1 as the genome of the experimentally tested, nicotine growth-negative *P. simplex* DSM 1323 (Figure 1b) was unavailable at the time of this writing. VPP protein sequences were obtained from accessions GCF_001652565.1 (*Shinella sp.* HZN7)^25^ and GCF_001551895.1 (*Agrobacterium tumefaciens* S33)^45^. Tblastn searches were done with BLAST 2.17.0+.

FastANI v1.34^22^ with default parameters was used to calculate pairwise average nucleotide identity (ANI) values. Digital DNA-DNA hybridization calculation was performed by submission of the *P. frigoritolerans* NIC8 chromosome to https://ggdc.dsmz.de/ggdc.php (accessed 10 August 2026) against reference NCBI accessions JALJWT000000000, CP017704, LGYA00000000 for *P. frigoritolerans* DSM 8801, *P. simplex* DSM 1321, and *P. butanolivorans* DSM 18926 respectively.

The cytochrome c HMMER search was carried out with HMMER 3.4^46^ using PF00034 as the model (https://www.ebi.ac.uk/interpro/entry/pfam/PF00034, accessed 10 August 2026).

For synteny comparisons of nicotine degrading pathways (Supporting Figure S2), genomes were downloaded from GenBank via accessions GCA_000219705.1 for *Pseudomonas putida S16*, GCF_001652565.1 for *Shinella sp.* HZN7, and GCF_021919345.1 for *Paenarthrobacter nicotinovorans* type strain ATCC 49919. Annotations of nicotine degradation pathway genes were manually cross referenced against enzymology, proteomics, and transcriptomics reports for *Pseudomonas putida S16*^12,26^, *Shinella sp.* HZN7^3,25,47^, and *Paenarthrobacter nicotinovorans*^24,48,49^ to ensure accuracy.

For phylogenetic tree analysis, the Ncox sequence was used a query to identify the top 5000 hits in a BLAST search run in August 2026. Sequences shorter than 350 amino acids were removed, and redundant sequences were removed using a 98% sequence identity threshold using Jalview 2.11.5.2. The resulting 3756 sequences were aligned using MAFFT v7.490 with the FFT-NS-2 strategy. The phylogenetic tree was then generated using IQ-TREE version 3.1.3 using the optimal substitution model (Q.PFAM+F4+G4) identified by the ModelFinder routine in IQ-TREE.

Multiple sequence alignments in Figure 2a were carried out with MAFFT^50^ v7.526 using flag -- auto and displayed in Jalview 2.11.5.0. Percent identities were computed using the NCBI BLAST webserver, accessed 01 December 2025. N-terminal signal sequences were identified with SignalP 6.0^51^ using the SignalP webserver (https://services.healthtech.dtu.dk/services/SignalP-6.0/, accessed 12 August 2026). Rossman folds were identified with Cofactory 1.0^52^ (https://services.healthtech.dtu.dk/services/Cofactory-1.0/, accessed 12 August 2026).

### Nicotine growth phenotyping

*P. frigoritolerans* DSM 8801 was obtained from Leibniz Institute DSMZ. *P. simplex* DSM 1323 was obtained from ATCC (ATCC 49098). A colony of *P. frigoritolerans* NIC8 that was *pnox+* was streaked onto an M9-traceLB-nicotine plate along with a *P. frigoritolerans* NIC8 *pnox-*(plasmid-cured) colony. For comparisons with other strains of *Peribacillus,* a colony of *P. frigoritolerans* NIC8, a colony of *P. frigoritolerans* DSM 8801, and a colony of *P. simplex* DSM 1323 were similarly streaked. Plates were incubated at 30° C and imaged after two days.

### RNA sequencing and analysis

Four colonies from an M9-traceLB-nicotine agarose plate were each inoculated into 3 mL M9-traceLB-glycerol (0.4% glycerol v/v) and grown overnight. Overnight cultures were diluted 1:100 into 3 mL M9-traceLB-glycerol or M9-traceLB-nicotine and grown for 10 hours with shaking at 200 rpm at 30° C. An OD_600_ value of 0.4 or 0.6 was reached for glycerol cultures or nicotine cultures respectively, as nicotine cultures appeared to grow slightly quicker. 1.5 mL from each culture was spun down, supernatant discarded, and pellet flash frozen in liquid nitrogen. A total of 8 pellets, 4 from each of nicotine or glycerol growth condition were shipped to GENEWIZ® NGS Services from Azenta Life Sciences (South Plainfield, NJ, USA) on dry ice for RNA extraction and sequencing as follows:

Total RNA was extracted using Qiagen Rneasy Plus Mini kit following manufacturer’s instructions (Qiagen, Hilden, Germany). Samples were then treated with TURBO DNase (Thermo Fisher Scientific, Waltham, MA, USA) to remove DNA contaminants. RNA samples were quantified using Qubit 4.0 Fluorometer (ThermoFisher Scientific, Waltham, MA, USA) and RNA integrity was checked with 4200 TapeStation (Agilent Technologies, Palo Alto, CA, USA). All 8 samples had a TapeStation RINe score > 9. RNA depletion sequencing library was prepared by using QIAGEN QIAseq FastSelect - 5S/16S/23S Kit (Qiagen, Hilden, Germany). RNA sequencing library preparation used the NEBNext Ultra II RNA Library Prep Kit for Illumina by following the manufacturer’s recommendations (NEB, Ipswich, MA, USA). Briefly, enriched RNAs were fragmented for 15 minutes at 94 °C. First strand and second strand cDNA were subsequently synthesized. cDNA fragments were end repaired and adenylated at 3’ends, and universal adapters were ligated to cDNA fragments, followed by index addition and library enrichment with limited cycle PCR. Sequencing libraries were validated using the Agilent Tapestation 4200 (Agilent Technologies, Palo Alto, CA, USA), and quantified using Qubit 4.0 Fluorometer (ThermoFisher Scientific, Waltham, MA, USA) as well as by quantitative PCR (KAPA Biosystems, Wilmington, MA, USA).

The sequencing libraries were multiplexed and clustered onto a flow cell on the Illumina NovaSeq instrument according to manufacturer’s instructions. The samples were sequenced using a 2x150bp Paired End (PE) configuration. Image analysis and base calling were conducted by the NovaSeq Control Software (NCS). Raw sequence data (.bcl files) generated from Illumina NovaSeq was converted into fastq files and de-multiplexed using Illumina bcl2fastq 2.20 software. One mis-match was allowed for index sequence identification. The number of reads per sample ranged from 28 million to 38 million. Additional statistics are reported in Supporting Table S6.

Fastq files were quality controlled using FastQC v. 0.12.1. Bowtie 2 v.2.5.4 was used to align reads to the full genomic sequence of *P. frigoritolerans* NIC8. featureCounts v2.0.3 was used to count reads for each gene with options -p --countReadPairs -T 8 -M -O -t gene -g locus_tag using the NCBI Prokaryotic Genome Annotation Pipeline (PGAP) gff3 file. DESeq2 v. 1.52.0 was used to compute differential expression statistics for glycerol vs. nicotine. Nicotine degrading operons were visualized with Integrative Genomics Viewer^53^ using the genome gff3 file as well as BigWig files generated from nicotine-grown samples’ .bam files using bamCoverage in DeepTools2^54^ using option -bs 10.

For verification of annotation robustness, Prokka v. 1.14.6 was used to re-annotate the contigs.fasta file using default options, and DESeq2 analysis showed no noticeable differences from the PGAP output.

Circular plots were made with differential expression results (Figure 4a, Supporting Figure S1) and visualized with pyCirclize (https://moshi4.github.io/pyCirclize/).

### Protein expression and steady state kinetics

*Ncox* and *pnox* were obtained as *E. coli* codon-optimized inserts in a pET28-his-SUMO-vector^19^ from Twist Biosciences. *E. coli* BL21 (DE3) was transformed with plasmids and colonies were grown overnight in LB at 37 °C. Resulting overnight cultures were diluted 100-fold into 3 L Protein Expression Medium (12 g L^−1^ tryptone, 24 g L^−1^ yeast extract, 50.4 g L^−1^ glycerol, 2.1 g L^−1^ K_2_HPO_4_ and 12.5 g L^−1^ KH_2_PO_4_) with 50 μg/mL kanamycin and grown to an OD_600_ of ∼1.0 at 37 °C. The flask was then shaken at 20 °C and expression was induced with 100 μM IPTG overnight. *E. coli* cells were pelleted by centrifugation, and cell pellets were stored at −20 °C. The pellet was resuspended in lysis buffer (50 mM Tris-HCl, 400 mM NaCl, 15 mM imidazole and 10% glycerol, pH 8.0) containing DNase I and cOmplete^TM^ protease inhibitor cocktail. The cell suspension was sonicated with a tip sonicator (Fisher Scientific Model 505) at 4 °C and solids removed by centrifugation twice at 30,000 x *g* and 4 °C for 30 min each time. The resulting supernatant was loaded onto a 5-mL HisTrap column pre-equilibrated with lysis buffer. The bound protein was first washed with 25 mL of lysis buffer and then 25 mL of lysis buffer supplemented to 10 mM imidazole, and protein was eluted in 10 mL lysis buffer with 500 mM imidazole. Eluted protein was dialyzed against 40 mM Tris-HCl, 0.2 M NaCl pH 8.0 in the presence of 40 μg of protease ULP-1 to cleave the His-SUMO tag. Dialyzed, cleaved protein was loaded onto the HisTrap column to remove the tag. The resulting eluate was diluted into 25 mM Tris-HCl pH 8.5 and loaded onto a 5 mL HiTrap Q column on an ӒKTA^TM^ Pure instrument (Cytiva) equilibrated in the same buffer. Protein was eluted using a linear salt gradient from 0 to 1 M NaCl. Purified protein was concentrated with a 30 kDa MWCO Amicon ® spin filter (Millipore Sigma) and stored at −80 °C until use (Supplementary Fig. S3).

All biochemical experiments were carried out in 40 mM HEPES, 100 mM NaCl at pH 7.4 with 10% (v/v) glycerol unless otherwise specified. The stock concentrations of Ncox and Pnox were determined by measuring the absorbance at maximum peak height between 445 and 460 nm on a Shimadzu UV-1900 UV-vis spectrophotometer, corresponding to the major flavin absorbance peak, using ε = 11300 M^-1^ cm^-1^.

Steady state kinetics analysis was carried out for Ncox at 20 °C by mixing 250 μL nicotine at varying concentrations with 250 μL enzyme at 20 nM and monitoring absorbance change at 280 nm on a Shimadzu UV-1900 UV-vis spectrophotometer, which corresponds to the major absorbance region of pseudooxynicotine that is not shared by nicotine. Linear plots were fitted to obtain the slope, and then converted to the correct units using the pseudooxynicotine-specific A_280_ extinction coefficient of 2914 M^-1^ cm^-1^ ^19^.

Steady state kinetics analysis was similarly carried out for Pnox at 20 °C by mixing 250 μL pseudooxynicotine (AmBeed, Inc. A285516) at varying concentrations with 250 μL enzyme at 200 nM and monitoring absorbance change at 288 nm on a Shimadzu UV-1900 UV-vis spectrophotometer, which corresponds to a wavelength of measurable absorbance difference between pseudooxynicotine and product. Linear plots were fitted to obtain the slope and converted to the correct units using the differential absorbance-specific A_288_ extinction coefficient of -1112 M^-1^ cm^-1^, derived from comparison of absorption spectra of pseudooxynicotine and pseudooxynicotine reacted to completion with Pnox.

Resulting nicotine or pseudooxynicotine concentration vs. enzyme velocity plots were fit to the Michaelis-Menten equation in GraphPad Prism v. 10.6.1.

To monitor flavin reduction upon substrate addition (Figures 5a, 5b), 75 μL of Ncox or Pnox at 20 μM was mixed with 75 μL 6 mM nicotine, pseudooxynicotine, or buffer alone, and the absorption spectrum was measured on a Shimadzu UV-1900 UV-vis spectrophotometer.

For the Amplex® Red assay, 40 μL of enzyme at 200 nM was mixed with 40 μL of substrate at 200 μM and the reaction was allowed to proceed for one hour at room temperature. 2 μL of the reaction was then diluted into 98 μL buffer and mixed with 100 μL Amplex® working solution (50 μM Amplex® UltraRed reagent, 0.2U/mL horseradish peroxidase). Fluorescence was quantified on a Tecan M1000 Infinite® Pro at an excitation wavelength of 565 nm and an emission wavelength of 597 nm with a 5 nm bandwidth for each.

### Transient kinetics

Transient kinetics experiments were conducted on a TgK Scientific SF-61DX2 KinetAsyst stopped-flow instrument.

Approximately 4 mL of 40 µM Ncox or Pnox, as determined by flavin absorbance, was placed in a glass tonometer and made anaerobic by cycling between argon and vacuum^55^. To measure flavin reduction with nicotine or pseudooxynicotine, a range of concentrations of the substrate in buffer was prepared anaerobically by argon sparging, and reactions were monitored by single-wavelength detection at 450 nm at 20 °C. The resulting curves were fit to a triexponential decay (equation 1) using custom R scripts in RStudio v. 2025.09.2 limiting analysis to after the first 0.002 s. Plots of the resulting observed rate constants versus substrate concentration were fit to a one-site binding model^31^ to determine substrate K_d_ values.

Equation 1: triple exponential decay. *A, B*, and *C* are pre-exponential weights for each exponential process defined by rates *k_obs,1_*, *k_obs,2_*, and *k_obs,3_*. *t* is time and *D* is baseline signal.

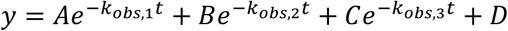

To measure the rate of oxygen reaction with reduced Ncox or Pnox, enzyme was reduced with 1 eq. nicotine or pseudooxynicotine in an anaerobic glass tonometer, with absorbance spectra obtained to confirm flavin reduction. For reactions of Ncox in the presence of 2 mM pseudooxynicotine, 440 μL of 20 mM pseudooxynicotine was added to a glass arm of the tonometer, made anaerobic at the same time as 4 mL of Ncox, and then mixed in after the enzyme had been reduced by 1 eq. of nicotine. Resulting anaerobic, reduced enzyme was reacted with buffer containing dissolved O_2_ at concentrations measured by a Hansatech Oxygraph+ liquid phase oxygen electrode system, with a range generated by sparging oxygen and nitrogen at different mixing ratios. The reaction was carried out at 20 °C and monitored using a multiwavelength CCD detector, and the resulting time vs. 450 nm curves were fit with a single exponential model for Ncox (equation 2) or a biexponential model for Pnox (equation 3) using a custom R script. The oxygen concentration vs. k_obs_ curves were fit in good agreement to lines at the oxygen concentrations tested to yield second order rate constants.

Equation 2: single exponential gain. *A* is the pre-exponential factor for the gain process defined by rate *k_obs_*. *t* is time and *B* is baseline signal.

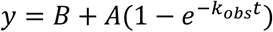

Equation 3: double exponential gain. *A,* and *B* are pre-exponential weights for each exponential gain process defined by rates *k_obs,1_* and *k_obs,2_*. *t* is time and *C* is baseline signal.

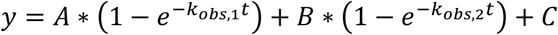

### Pseudomonas expression and growth curves with Ncox and Pnox

*Ncox* and *pnox* were obtained as gBlocks (IDT) with flanking EcoRI and XbaI sites with codon optimization for *Pseudomonas putida*. Gene fragments were digested with EcoRI and XbaI and ligated into the correspondingly digested pJN105-NicA2 vector. Resulting constructs were transformed into DH10B *E. coli* cells with gentamicin selection and sequence-confirmed through whole plasmid sequencing at Plasmidsaurus. pJN105-Ncox and pJN105-Pnox were transformed into electrocompetent *P. putida* S16 Δ*cycN*.

Δ*cycN* strains were grown for 24 hours in M9-glycerol-gentamicin (6 g/L Na_2_HPO_4_, 3 g/L KH_2_PO_4_, 1 g/L ammonium chloride, 1 mM MgSO_4_, 0.1 mM CaCl_2_, 1× trace metals (Teknova, T1001), 1 μg/mL thiamine, 0.4% v/v glycerol, 25 μg/mL gentamicin) at 30 °C. Resulting cultures were washed once with M9 salts (6 g/L Na_2_HPO_4_, 3 g/L KH_2_PO_4_, 1 g/L ammonium chloride) and normalized to OD600 of 1.0. 2.5 μL of each culture was diluted into 250 μL M9-nicotine-gentamicin in a 96 well plate, with varying arabinose concentrations and a total of 3 technical replicates. A Breathe-Easy^TM^ sealing membrane plate cover (Sigma) was used to decrease evaporation. The plate was monitored with shaking at 750 nm (to minimize the impact of the colored pigment) for 7 days at 30 °C with data collection every 30 minutes.

For comparisons of protein expression in *P. putida* S16 Δ*cycN*, 30 μL of overnight-grown bacteria at 30 °C in M9-glycerol-gentamicin was added to 1.5 mL M9-nicotine-gentamicin containing 0.2% w/v arabinose and grown at 30 °C for 24 hours. Cultures were harvested and OD_600_-normalized, and 0.18 OD-mL of each sample pellet was resuspended in 50 μL M9, mixed with 25 μL reducing buffer (50% v/v glycerol, 10% w/v sodium dodecyl sulfate, 0.0005% w/v bromophenol blue, 5% v/v 2-mercaptoethanol, 300 mM Tris pH 7.0) and incubated at 95 °C for 10 minutes. 10 μL of each sample was loaded onto a 4-15% TGX mini-PROTEAN gel (Bio-Rad). Protein was visualized by a modification of the original Fairbanks Coomassie staining procedure^56^.

## Data availability

All strains are available upon request to James Bardwell. The genome sequence of *P. frigoritolerans* NIC8 is available at Genbank accession SAMN53626221. RNA sequencing data are available on the NCBI Gene Expression Omnibus under accession GSE344791.

## Supporting information

This article contains supporting information.

## Acknowledgements

We thank Ke Wan for protein purification assistance. This research was supported in part through computational resources and services provided by Advanced Research Computing at the University of Michigan, Ann Arbor. Research reported in this publication was supported by the University of Michigan Natural Products Discovery Core (NPDC).

## Funding and additional information

This work was funded by Howard Hughes Medical Institute where JCAB is an Investigator. This research was also supported by National Science Foundation grant 2236541 to FS. The NPDC is grateful for support from the U-M Life Sciences Institute and the U-M Biosciences Initiative (RRID: SCR_023105).

