## Supporting data file for "Isolation of oxygen-dependent nicotine- and pseudooxynicotine-metabolizing enzymes"

*Running title: Genuine nicotine and pseudooxynicotine oxidases*

*Keywords: oxidation-reduction (redox), flavoprotein, enzyme kinetics, bacterial genomics, transcriptomics*

Tejas A. Navaratna[1], Javeria Akram[2], Thea D. Pazdernik[1], Amudha Ramachandran[3], Pamela Schultz[3], Mark Dulchavsky[1], Xavier Choussat[1], Claire Oczon[1], Aashnaa Singh[1], Nikhil Myers[1], Aaron Robida[4], Ashootosh Tripathi[3], Frederick Stull[2], James C. A. Bardwell[1*]

1. Howard Hughes Medical Institute and Department of Molecular, Cellular and Developmental Biology, University of Michigan, Ann Arbor, MI, USA
2. Department of Chemistry, Western Michigan University, Kalamazoo, MI, USA
3. Natural Products Discovery Core, Life Sciences Institute, University of Michigan, Ann Arbor, MI, USA
4. Center for Chemical Genomics, Life Sciences Institute, University of Michigan, Ann Arbor, MI, USA

# S1

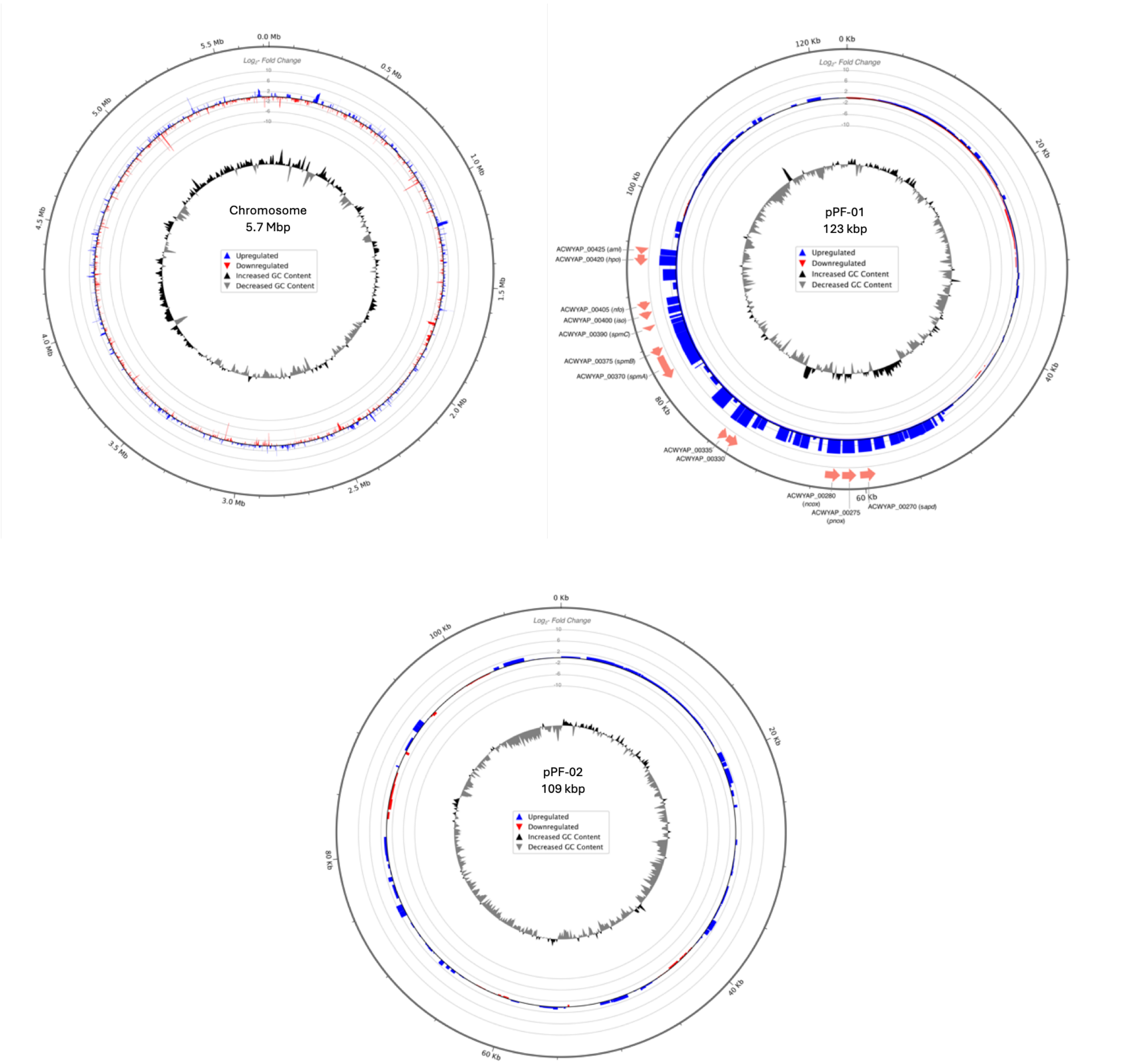

**Figure S1** – representations of the chromosome and plasmids of *Peribacillus frigoritolerans var. nicotinobibens.* Also depicted are RNAseq $log_2$fold-change values for open reading frames and GC content. The nicotine-degrading catabolic island is apparent in the bottom-left quadrant of plasmid pPF-01.

# S2

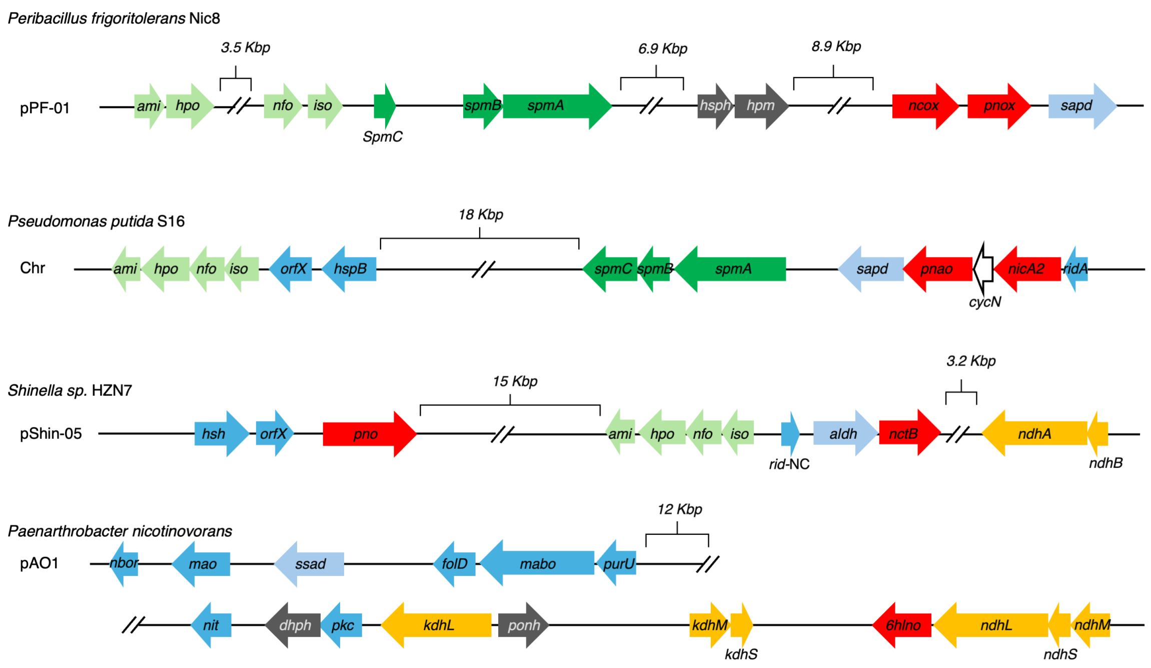

**Figure S2** – enzymes with direct or proposed enzymatic roles in *Peribacillus frigoritolerans* NIC8 (this work), *Pseudomonas putida* S16's pyrrolidine pathway, *Shinella sp.* HZN7's VPP pathway, and *Paenarthrobacter nicotinovorans*' pyridine pathway of nicotine metabolism. Note that *P. nicotinovorans* diagram has been condensed to two lines for brevity. Open reading frames involved in gene regulation (e.g. transcription factors), transposon elements, and other open reading frames are not shown. Of note, *P. putida* S16 contains an open reading frame between *nicA2* and *pnao* encoding a cytochrome C. A cytochrome C homolog is not present in the pPF-01 plasmid of *P. frigoritolerans* NIC8.

# S3

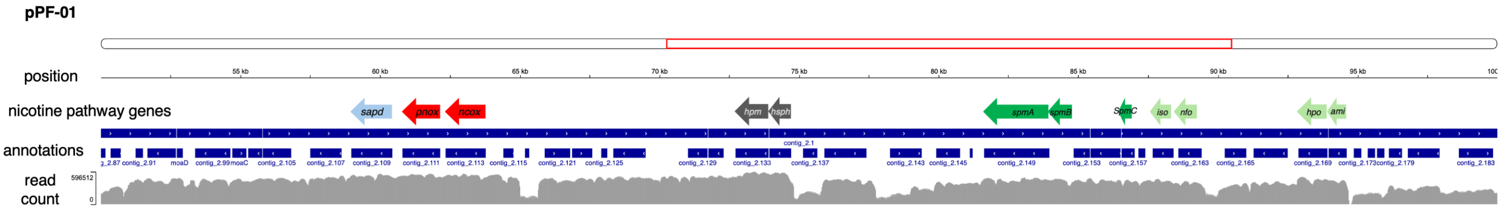

**Figure S3** – RNA sequencing shows that nicotine degrading pathway genes of *Peribacillus frigoritolerans* N8 are localized in plasmid 1 on four operons: (1) operon with *ncox, pnox,* and *sapd*; (2) operon with *hsph* and *hpm*; (3) operon with *nfo, iso, and spmABC*; and (4) operon with *ami* and *hpo*. This figure was generated with Integrative Genomics Viewer using the genome .gff3 file and RNA sequencing data of nicotine-grown samples.

# S4

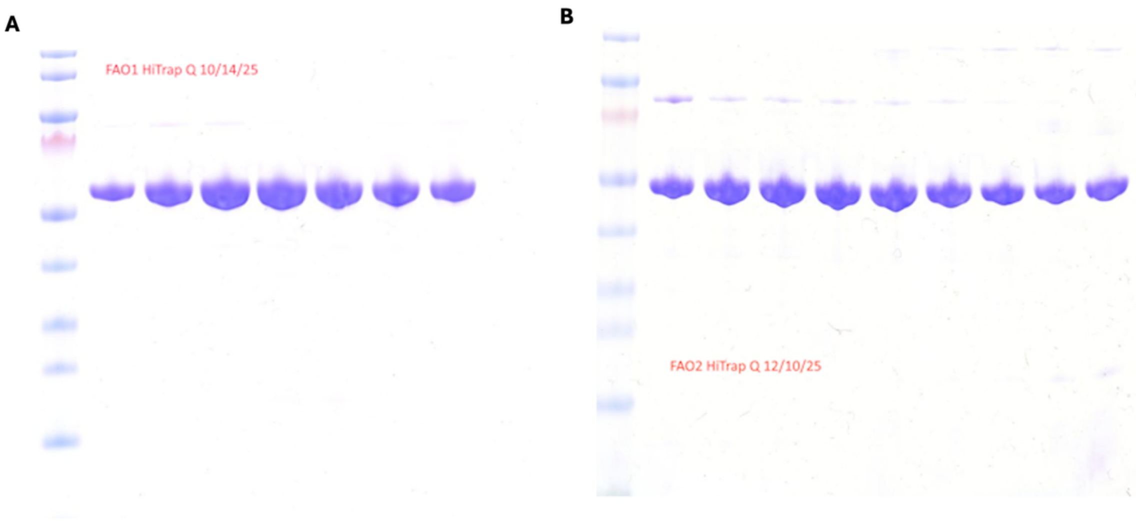

**Figure S4** –**a,** Coomassie blue-stained protein gel of Ncox; all fractions corresponding to lanes 1-7 were combined following cation exchange chromatography with HiTrap Q. **b,** Coomassie blue-stained protein gel of Pnox. Fractions corresponding to lanes 2-8 were combined following cation exchange chromatography with HiTrap Q.

# S5

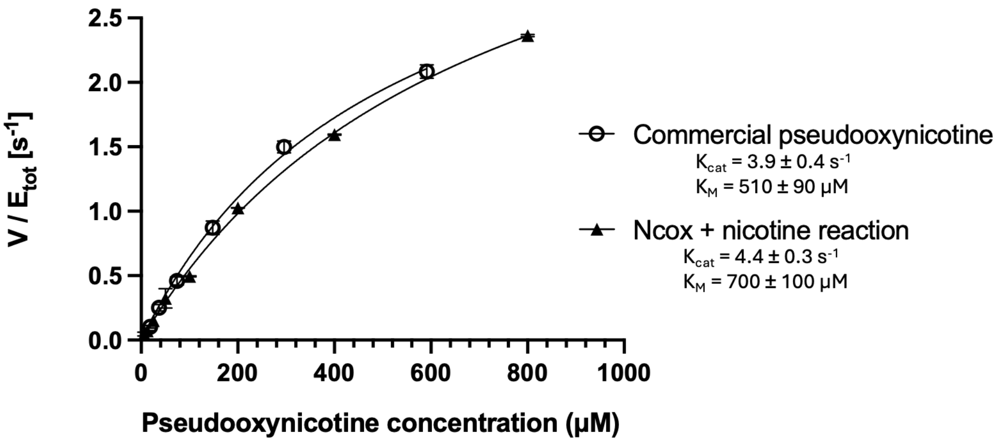

**Figure S5** – Michaelis-Menten kinetics analysis of Pnox with commercially-obtained pseudooxynicotine or pseudooxynicotine generated by reaction between Ncox and nicotine.

# S6

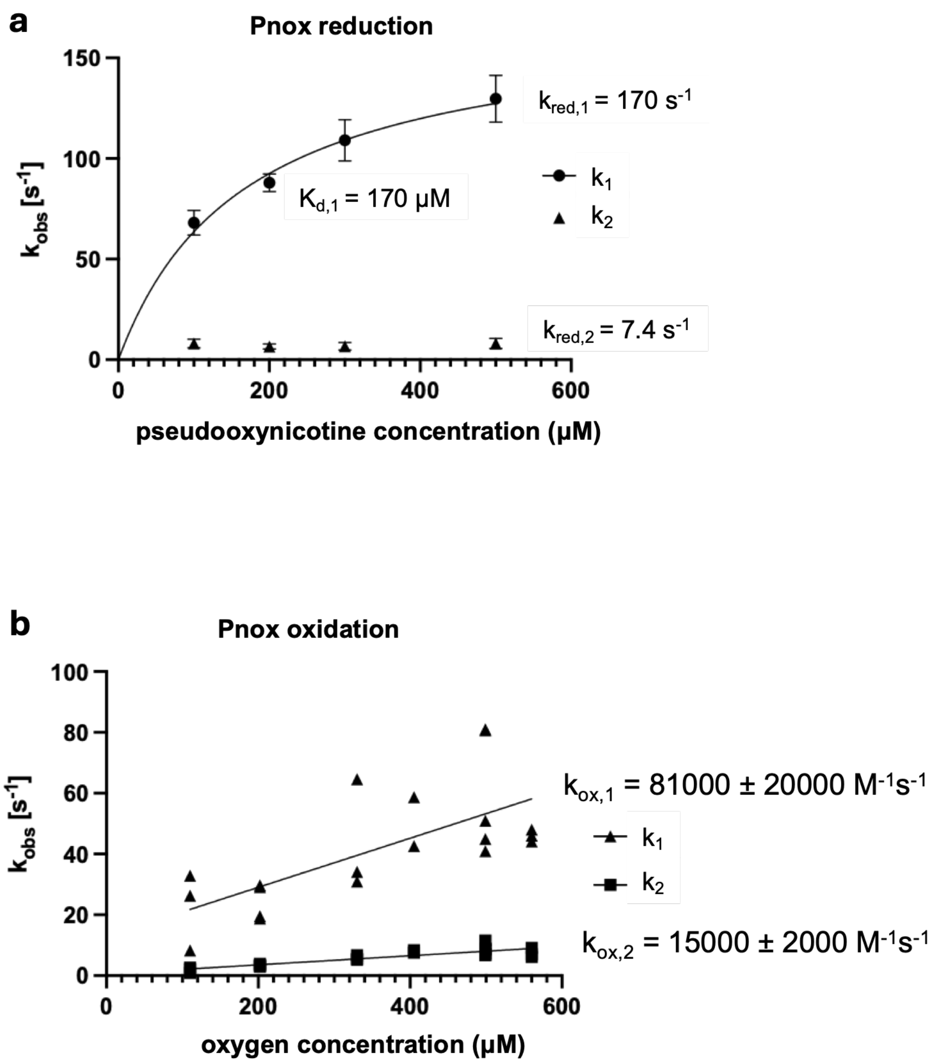

**Figure S6** – Pnox half-reaction data showing **a**, reduction half reaction rate constant 1, $k_{obs,1}$, shows a monotonic dependence on pseudooxynicotine concentration that can be fit to a binding isotherm, yielding a $K_d$ of 170 μM. **b**, oxidation half reaction is best-fit with a biexponential equation, and shows a linear relationship with oxygen concentration, with the second-order rate constants of 81000 M$^{-1}$s$^{-1}$ and 15000 M$^{-1}$s$^{-1}$.

# S7

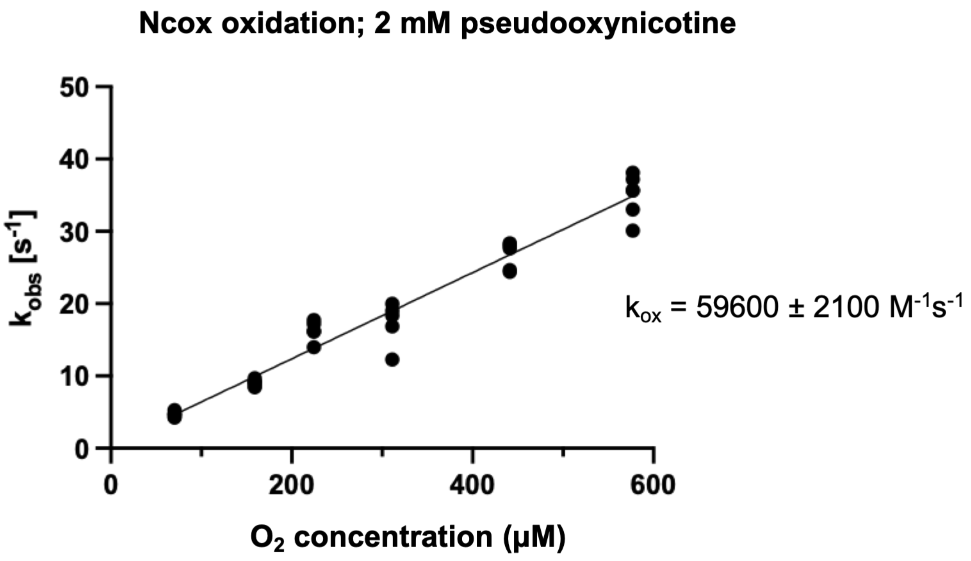

**Figure S7** - Ncox oxidation half-reaction is unaffected by the presence of 2 mM pseudooxynicotine, the product of nicotine oxidation.

# S8

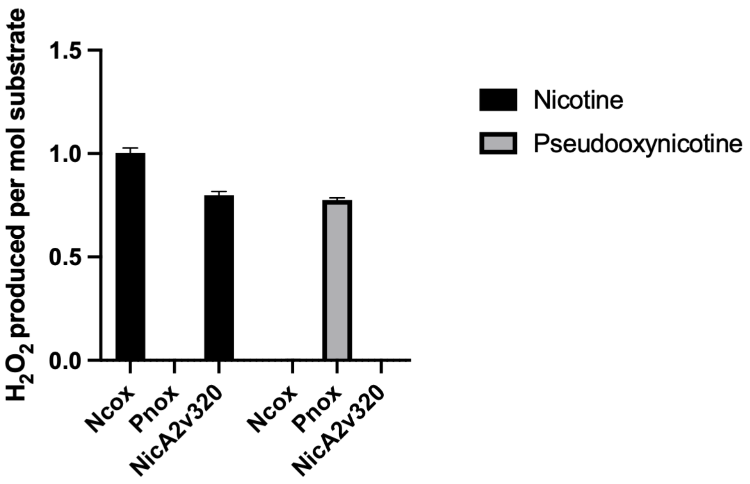

**Figure S8** – Measurement of $H_2O_2$ production for enzymes Ncox, Pnox, and NicA2v320 for the substrates nicotine and pseudooxynicotine. Enzymes at 100 nM were incubated with 100 μM substrate for one hour at room temperature.

## S9

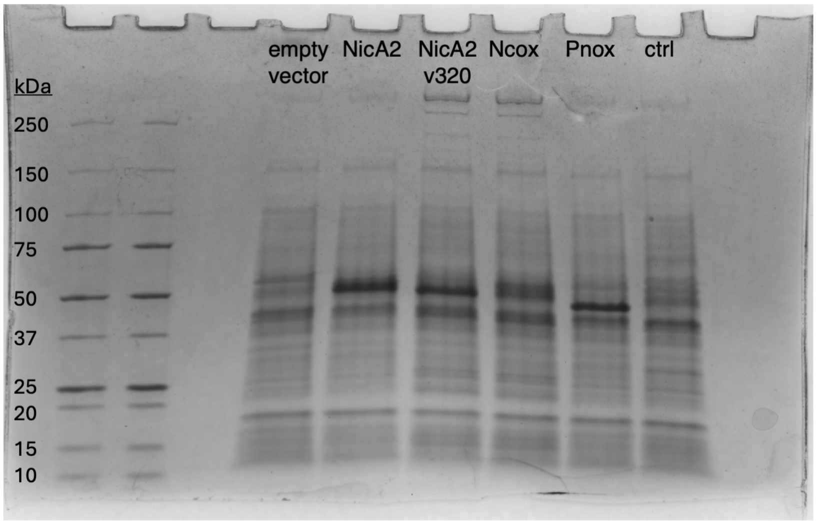

**Figure S9** – whole cell SDS-PAGE of *Pseudomonas putida* S16 Δ*cycN* with pJN105 plasmids expressing empty vector, NicA2, NicA2v320, Ncox, Pnox, or a low-expressing control protein. Each strain was grown in M9 nicotine with 0.2% w/v arabinose for 24 hours following 1:50 dilution from an overnight M9 glycerol culture.

**Table S1** – DNA sequences used in this work. All are shown in 5’ to 3’ direction.

| Name | Sequence |
|---|---|
| 16S amplicon | GGCAACCTGCCTATAAGACTGGGATAACTTCGGGAAACCGGAN<br>CTAATACCGGATACGTTCTTTTCTCGCATGAGAGAAGATGGAAA<br>GACGGTTTACGCTGTCACTTATAGATGGGCCCGCGGCGCATTA<br>GCTAGTTGGTGAGGTAATGGCTCACCAAGGCGACGATGCGTAG<br>CCGACCTGAGAGGGTGATCGGCCACACTGGGACTGAGACACG<br>GCCCAGACTCCTACGGGAGGCAGCAGTAGGGAATCTTCCGCA<br>ATGGACGAAAGTCTGACGGAGCAACGCCGCGTGAACGAAGAA<br>GGCCTTCGGGTCGTAAAGTTCTGTTGTTAGGGAAGAACAAGTA<br>CCAGAGTAACTGCTGGTACCTTGACGGTACCTAACCAGAAAGC<br>CACGGCTAACTACGTGCCAGCAGCCGCGGTAATACGTAGGTGG<br>CAAGCGTTGTCCGGAATTATTGGGCGTAAAGCGCGCGCAGGTG<br>GTTCCTTAAGTCTGATGTGAAAGCCCACGGCTCAACCGTGGAG<br>GGTCATTGGAAACTGGGGAACTTGAGTGCAGAAGAGGAAAGT<br>GGAATTCCAAGTGTAGCGGTGAAATGCGTAGAGATTTGGAGGA<br>ACACCAGTGGCGAAGGCGACTTTCTGGTCTGTAACTGACACTG<br>AGGCGCGAAAGCGTGGGGAGCAAACAGGATTAGATACCCTGG<br>TAGTCCACGCCGTAAACGATGAGTGCTAAGTGTTAGAGGGTTTC<br>CGCCCTTTAGTGCTGCAGCTAACGCATTAAGCACTCCGCCTGG<br>GGAGTACGGCCGCAAGGCTGAAACTCAAAGGAATTGACGGGG<br>GCCCGCACAAGCGGTGGAGCATGTGGTTTAATTCGAAGCAACG<br>CGAAGAACCTTACCAGGTCTTGACATCCTCTGACAACCCTAGA<br>GATAGGGCTTTCCCCTTCGGGGGA |
| Ncox insert for growth curves | gcgaattcaggaggatatcaccatgaataaccgccagaacaaaacccaaaacttcgacgt<br>ggtcatcatcggtggcggcttcaccggtgtcaccgccgcacgtgagctccgccataagggct<br>acagcgtggtgctgctggaggcgaaagaccacctcggcgggcgtacctggttcgacgagc<br>gcttgggcaccgaagtggagctcggtggcacctgggtgcactccgtgcagccccatacgtg<br>gtccgaaatctcccggtacggcctggaactcttggactacagcaacatcgaacaccaggcc<br>tactggattgccgatggcaagagcaacagcggcagcatggccgatctggctggcctgctgg<br>acaagggcatgaacctgtcgttcgagggctgccgcgaatactttccgaacccgtacgaccc<br>actgctgagcccgagcatctcggaaattgatggcatgagcttcgcagaccggttgaagtcca<br>tggatctcaccaaggaggagtacgatgtggtggagggcatctgggacctgaatttctcgtcgt<br>cgctggaagatggcggtctgtcggcggccctgcgttggggctcgctctgtcgcggcgactgg<br>cacgtgttcttcgagaccagcaccctctaccgcctgaagaagggcacccgcagcctgtaca<br>cggctatcgccgaagacggcggcgccgaaattcgcctgagcaccgctgtgtcggccatcg<br>agcgcacggacaatggtgtgattgtgcgtacccgcgacggccaacagatccaagccaaag<br>cagtcgtggtggcggccccccctgaacagcctgaccaacatcgagttcaccccggccctgtc<br>cgaggtcaagcaggccgccttggccgagggctacccatcgcgcggcgtcaagttcatgatg<br>cgtttgaaaggtaagctggatccgtttgtggctgtcgccccggccggtagcaagctgagctat<br>gtccacctgaacggctacgtcgacgacgactccattgcggtcgccttcgggagcgacgtgg<br>cccgcatcgatatgaacgacgtgagcgcggtggaggcggagctgcgtaagtggattccgg<br>acatcgaggtggtcgccaccgctgggttcgattacaagcgcgatgagttcaccaatgagac<br>gtggagcatgttccgtccgaacatgctgacccgctacttgaaagagctgcagcgccctgaaa<br>atggcgtcttcctggccggctcggacatcgccagcggctggaccggttatatcgacggcgcg<br>atcgagtcgggcctgaccgccggtcgtaacgtgcacgaatacatcaccaatagcagcaag |

| | |
|---|---|
| | gacgtcatcgtcttcgacaacatcgatggtgtcatcgagaacaactccaccgcgagcaaca acgtccaggagtacatcatcatcccttccgaggacgtggtggccccgccgaagtaatctaga gcggc |
| Pnox insert for growth curves | gcgaattcaggaggatatcaccatgcaggtgaacaacaccgaaaagaacaactacgatgt catcatcattggcgcgggcttcgcgggcgtgacggcggcccgtgaactgcgccgcatgggt cggaccgcgctgatcgtggaagcacgcgatcggctgggcggtcgtacctggaccgaagat cgtctgggggcccagctggaaatcggcggcacctgggtccaccccatccagccgaacgtgt gggccgaaattatgcggtacggtctggagctgatttcgtcgccagcccagaagtatgcccact ggattgccgacggccaactcaagagcggctcgatcgaggaatttgcaaagctggtggaca acgcgtataaccgggtcctggaggatagccggttccacctgtacaacccctacgacccgctc tcgagcgaaaccctggaggaaatggacaaacaatcgttcaccgaccgcctggacagcctg gacctgaccaaggaggaatacgacctgatgcatggcatgtgggccaccagcttccaggcg ccgccggaagaaggcggcctgagcagcgctatccgctggggcgctttgtcctgggggtcct ggcagctcatgctggagatgctgtgcgtgtacaagctgaagaaggggacccgcgccctgat tgaagcaatggccgcggacgctgccgcagacaccaaattctcgaccatcgtgacgtcgatt gagaaaaccgacgacggtgtcaccgtgtacaccaaggacggccaacagctccagggca aagccgtcatcattaccgtgccactgaatgtgctgaagtcgatcgagttcacccctccgctgtc cgagggtaaaatgatcgtcgccaacgaagggcaagccagccggggtgtcaaggtgatcg cccgcatccgtggcgagttcgagccatttatggcggctgcccccggcaactatcccctggcat acgcgcagctggaataccacgtcgagggtgatagcatcgtcgtggctttcggcgcggatgc aaccaagctggacgcaaacgatgtcagcgcggtgtcgaatgccttccgccagtggctgccg gacgtcgaagtcgtcgcgagcacgggccacgactgggtcgccgatgaatttagccaagaa acctggtacatgtcgcggccgaaccagctgcgctacctgggtgaactgcagcgcccggaa aacggcatgtttttggccggctcggactacgccagcggttgggctggcttcattgatggggcc atcgaatcgggcatgttggtcagccgcaacgtccacgagtacatcacctcgaacgacgcgc agaccaacgacgtcgtcggcacccacaagtaatctagagcggc |
| His-SUMO-Ncox insert for purification | CCCCTCTAGAAATAATTTTGTTTAACTTTAAGAAGGAGATATACCA TGGGCAGCAGCCACCACCACCATCATCATCACCATCATCACAG CAGCGGCCTGGTGCCGCGCGGCAGCCATATGGCTAGCATGTC GGACTCAGAAGTCAATCAAGAAGCTAAGCCAGAGGTCAAGCCA GAAGTCAAGCCTGAGACTCACATCAATTTAAAGGTGTCCGATG GATCTTCAGAGATCTTCTTCAAGATCAAAAAGACCACTCCTTTAA GAAGGCTGATGGAAGCGTTCGCTAAAAGACAGGGTAAGGAAAT GGACTCCTTAAGATTCTTGTACGACGGTATTAGAATTCAAGCTG ATCAGACCCCTGAAGATTTGGACATGGAGGATAACGATATTATTG AGGCTCACAGAGAACAGATTGGTGGATCCATGAACAACCGACA GAACAAGACTCAGAATTTTGACGTCGTCATAATAGGCGGCGGC TTCACCGGTGTCACAGCTGCACGAGAGCTCCGACACAAAGGTT ACTCCGTTGTACTTCTTGAGGCTAAAGATCACTTAGGCGGTCGG ACGTGGTTCGACGAGAGACTTGGAACCGAGGTTGAGCTGGGT GGTACCTGGGTTCACTCGGTTCAACCCCACACTTGGTCAGAGA TCTCCCGGTACGGATTGGAGTTACTGGATTATTCTAACATCGAA CACCAAGCGTACTGGATCGCAGACGGTAAGTCAAACTCGGGGA GCATGGCTGATTTGGCTGGCTTGCTGGACAAAGGCATGAACCT GTCATTCGAAGGATGTCGGGAGTACTTCCCGAACCCTTACGAC CCACTGCTTTCACCCTCTATAAGCGAGATAGATGGCATGAGTTT |

| | |
|---|---|
| | CGCCGATCGGCTTAAGAGTATGGACCTTACAAAAGAAGAGTATG ACGTTGTTGAGGGAATATGGGACTTAAATTTCTCTTCCTCATTAG AAGACGGTGGATTATCGGCCGCGCTCCGTTGGGGCAGCCTGT GTCGAGGGGACTGGCATGTTTTCTTTGAGACCTCAACCCTCTA CCGGCTTAAGAAAGGTACGCGATCGCTCTATACGGCGATTGCT GAAGATGGTGGCGCGGAGATTCGGTTATCAACCGCCGTCAGC GCGATCGAGCGGACAGATAATGGGGTGATCGTAAGAACGCGC GACGGCCAACAAATCCAGGCGAAGGCCGTAGTAGTTGCTGCA CCGCTTAACTCACTTACGAATATTGAATTTACTCCGGCGCTGAG CGAGGTTAAGCAGGCAGCTCTGGCGGAAGGTTACCCCAGCCG CGGCGTGAAGTTTATGATGCGTTTGAAGGGTAAGTTGGATCCCT TTGTCGCTGTAGCACCGGCAGGTAGCAAGCTTTCGTACGTGCA TTTAAACGGTTACGTAGACGACGACTCCATAGCCGTAGCCTTTG GCAGCGACGTAGCGCGCATCGACATGAACGACGTTTCAGCGG TAGAGGCGGAACTTCGTAAATGGATTCCGGATATCGAAGTTGTT GCTACCGCGGGCTTCGACTACAAGCGCGACGAGTTCACCAAC GAGACGTGGTCTATGTTTCGGCCGAACATGTTAACGCGTTACCT CAAAGAGTTACAACGCCCTGAAAATGGGGTCTTCTTGGCAGGT AGCGACATTGCAAGTGGATGGACCGGATATATAGACGGGGCTAT AGAGAGCGGCTTAACAGCGGGTCGTAACGTACATGAGTATATTA CTAATTCCAGTAAGGATGTAATAGTTTTCGACAACATTGACGGC GTGATCGAGAACAATAGCACAGCATCCAATAACGTCCAAGAATA TATTATTATCCCGAGCGAAGACGTTGTTGCGCCACCCAAGTGAT AACTCGAGCACC |
| His-SUMO-Pnox insert for purification | CCCCTCTAGAAATAATTTTGTTTAACTTTAAGAAGGAGATATACCA TGGGCAGCAGCCACCACCACCATCATCATCACCATCATCACAG CAGCGGCCTGGTGCCGCGCGGCAGCCATATGGCTAGCATGTC GGACTCAGAAGTCAATCAAGAAGCTAAGCCAGAGGTCAAGCCA GAAGTCAAGCCTGAGACTCACATCAATTTAAAGGTGTCCGATG GATCTTCAGAGATCTTCTTCAAGATCAAAAAGACCACTCCTTTAA GAAGGCTGATGGAAGCGTTCGCTAAAAGACAGGGTAAGGAAAT GGACTCCTTAAGATTCTTGTACGACGGTATTAGAATTCAAGCTG ATCAGACCCCTGAAGATTTGGACATGGAGGATAACGATATTATTG AGGCTCACAGAGAACAGATTGGTGGATCCATGCAAGTTAACAA CACTGAAAAGAATAACTACGACGTCATCATAATTGGAGCGGGAT TTGCTGGTGTGACTGCAGCACGGGAGCTGAGACGCATGGGTC GTACAGCCCTGATCGTGGAAGCGCGTGACCGCCTTGGTGGCA GAACCTGGACAGAAGACCGCCTGGGAGCACAACTTGAGATAG GTGGTACTTGGGTTCATCCGATTCAGCCGAATGTGTGGGCCGA GATAATGCGTTACGGTCTTGAATTAATTTCGAGCCCCGCTCAGA AGTACGCGCACTGGATCGCAGATGGTCAACTCAAATCAGGTAG CATAGAAGAGTTCGCTAAGCTGGTTGACAACGCTTACAACCGC GTTCTTGAGGACAGTCGCTTTCATTTGTATAACCCTTATGACCC GCTGTCTTCAGAGACATTAGAAGAGATGGACAAGCAGAGTTTC ACTGACCGCCTTGACTCATTAGATCTGACGAAGGAAGAGTATGA CTTGATGCATGGTATGTGGGCTACGTCCTTCCAGGCCCCACCG |

| | |
|---|---|
| | GAAGAGGGTGGGCTTTCAAGCGCAATACGTTGGGGCGCGTTG<br>TCGTGGGGATCATGGCAGCTTATGTTAGAAATGCTGTGTGTGTA<br>TAAGCTGAAGAAGGGCACTCGTGCGTTGATTGAGGCCATGGCG<br>GCGGATGCCGCGGCCGACACCAAGTTCAGTACAATTGTCACCA<br>GTATAGAAAAGACAGACGACGGTGTTACAGTTTACACGAAGGA<br>CGGCCAACAGTTACAGGGCAAGGCGGTGATCATAACTGTCCCT<br>CTGAACGTTCTGAAAAGTATTGAATTTACACCCCCACTGAGTGA<br>GGGGAAAATGATTGTGGCGAACGAGGGCCAAGCTAGCAGAGG<br>AGTTAAGGTGATTGCCCGCATTCGCGGGGAGTTTGAGCCTTTC<br>ATGGCCGCGGCCCCGGGGAATTACCCCCTGGCTTATGCACAAC<br>TGGAATACCATGTTGAGGGCGATAGTATTGTGGTTGCTTTCGGC<br>GCCGATGCAACGAAGCTCGATGCGAATGACGTCTCCGCAGTTA<br>GTAACGCGTTTAGACAGTGGTTACCTGACGTAGAGGTAGTTGCT<br>TCAACCGGTCATGACTGGGTGGCCGACGAGTTCTCTCAGGAAA<br>CATGGTACATGTCCCGGCCTAACCAATTACGCTACCTTGGGGAA<br>CTGCAGCGCCCTGAGAACGGAATGTTCCTGGCAGGTTCAGATT<br>ACGCAAGTGGTTGGGCTGGTTTCATTGATGGCGCTATCGAGAG<br>TGGCATGCTGGTTAGCCGAAACGTCCATGAGTACATTACTTCAA<br>ACGACGCGCAGACCAACGACGTCGTCGGAACCCACAAATAATA<br>ACTCGAGCACCACC |

**Table S2**: Genome sequencing statistics

| Contig name | Length (bp) | Coverage | Number of annotated genes | %GC |
|---|---|---|---|---|
| chromosome | 5722729 | 180 x | 5669 | 40.7 |
| Plasmid 1 (pPF-01) | 123363 | 341 x | 111 | 37.8 |
| Plasmid 2 (pPF-02) | 109425 | 303 x | 129 | 36.2 |

**Table S3** – homology to known nicotine degrading enzymes. Function abbreviations: 3-SSP, 3-succinoylsemialdehyde pyridine; HSP, 6-hydroxy-3-succinoyl pyridine; HP, 2-hydroxypyridine; 2,5-DHP, 2-5-dihydroxypyridine.

| Gene | Length | Function | Homolog | Organism | % identity | % coverage | Accession |
|---|---|---|---|---|---|---|---|
| ACWYAP_00280 (*ncox*) | 479 | Nicotine oxidoreductase | Pnao | *Pseudomonas putida* S16 | 37 | 89 | AEJ14619.1 |
| | | | NicA2 | *Pseudomonas putida* S16 | 35 | 86 | AEJ14620.1 |
| ACWYAP_00275 (*pnox*) | 449 | Pseudooxynicotine oxidase | Pnao | *Pseudomonas putida* S16 | 41 | 94 | AEJ14619.1 |

| | | | | | | | |
|---|---|---|---|---|---|---|---|
| | | | NicA2 | *Pseudomonas putida* S16 | 33 | 91 | AEJ14620.1 |
| ACWYAP_00270 (*sapd*) | 490 | 3-SSP dehydrogenase | Sapd | *Pseudomonas putida* S16 | 38 | 97 | AEJ14618.1 |
| | | | Aldh | *Shinella sp.* HZN7 | 35 | 93 | AGS16699.2 |
| ACWYAP_00370 (*spmA*) | 778 | 3-succinoylpyridine monooxygenase subunit A, molybdopterin-binding | NdhL | *Paenarthrobacter nicotinovorans* | 28 | 97 | AAK64263.1 |
| | | | KdhL | *Paenarthrobacter nicotinovorans* | 25 | 97 | AAK64253.1 |
| | | | SpmA | *Pseudomonas putida* S16 | 27 | 96 | AEJ14617.1 |
| ACWYAP_00375 (*spmB*) | 292 | 3-succinoylpyridine monooxygenase subunit B, FAD-binding | NdhM | *Paenarthrobacter nicotinovorans* | 29 | 99 | AAK64243.1 |
| | | | KdhM | *Paenarthrobacter nicotinovorans* | 28 | 99 | AAK64248.1 |
| | | | SpmB | *Pseudomonas putida* S16 | 30 | 43 | WP_080563817.1 |
| ACWYAP_00390 (*spmC*) | 156 | 3-succinoylpyridine monooxygenase subunit C, iron sulfur cluster-binding | NdhS | *Paenarthrobacter nicotinovorans* | 42 | 95 | AAK64244.1 |
| | | | KdhS | *Paenarthrobacter nicotinovorans* | 37 | 94 | AAK64247.1 |
| | | | VPPA$_S$/NdhS | *Shinella sp.* HZN7 | 42 | 88 | AMB57024.1 |
| | | | SpmC | *Pseudomonas putida* S16 | 36 | 92 | AEJ14616.1 |
| ACWYAP_00335 | 259 | Putative HSP hydrolase | | | | | |
| ACWYAP_00330 | 394 | Putative 2-HP hydroxylase | | | | | |
| ACWYAP_00420 (*hpo*) | 345 | 2,5-DHP dioxygenase | VppE/Hpo | *Shinella sp.* HZN7 | 53 | 98 | ANH08438.1 |
| | | | Hpo | *Pseudomonas putida* S16 | 53 | 95 | AEJ14599.1 |
| ACWYAP_00405 (*nfo*) | 278 | N-formylmaleamate deformylase | VppF/Nfo | *Shinella sp.* HZN7 | 31 | 86 | ANH08439.1 |
| | | | Nfo | *Pseudomonas putida* S16 | 32 | 84 | AEJ14600.1 |
| ACWYAP_00425 (*ami*) | 216 | Maleamate amidohydrolase | VppG/Ami | *Shinella sp.* HZN7 | 43 | 89 | ANH08437.1 |
| | | | Ami | *Pseudomonas putida* S16 | 41 | 94 | AEJ14598.1 |
| ACWYAP_00400 (*iso*) | 249 | Maleate-formate isomerase | VppH/Iso | *Shinella sp.* HZN7 | 53 | 99 | ANH08440.1 |
| | | | Iso | *Pseudomonas putida* S16 | 55 | 99 | AEJ14601.1 |

**Table S4** - top 20 upregulated genes. Numbers in experimental columns represent transcripts per kilobase million (tpm) counts. Log2Fc = $Log_2$(Fold-change) between nicotine and glycerol growth conditions.

| Locus tag | Annotation | Nic-1 | Nic-2 | Nic-3 | Nic-4 | Gly-1 | Gly-2 | Gly-3 | Gly-4 | Log$_2$Fc |
|---|---|---|---|---|---|---|---|---|---|---|
| 26215 | AEC family transporter | 568 | 474 | 478 | 447 | 6 | 6 | 5 | 5 | 6.93 |
| 00335 | alpha/beta fold hydrolase | 25205 | 24482 | 22778 | 23490 | 426 | 446 | 412 | 483 | 6.22 |
| 00330 | LLM class flavin-dependent oxidoreductase | 27540 | 27800 | 26338 | 27244 | 518 | 529 | 486 | 567 | 6.14 |
| 00420 | 2,5-dihydroxypyridine 5,6-dioxygenase NicX family protein | 2530 | 2902 | 2810 | 2682 | 55 | 52 | 53 | 58 | 6.11 |
| 00425 | isochorismatase family protein | 1466 | 1328 | 1334 | 1184 | 30 | 30 | 32 | 36 | 5.88 |
| 00345 | sodium:solute symporter family protein | 1435 | 1592 | 1602 | 1565 | 36 | 40 | 38 | 45 | 5.73 |
| 08410 | GapA-binding peptide SR1P | 316 | 657 | 657 | 810 | 9 | 13 | 18 | 18 | 5.71 |
| 00350 | DUF3311 domain-containing protein | 1363 | 1488 | 1476 | 1430 | 40 | 39 | 36 | 51 | 5.57 |
| 02435 | BH0509 family protein | 294 | 315 | 345 | 253 | 12 | 12 | 14 | 15 | 5.03 |
| 00280 | flavin monoamine oxidase family protein | 9129 | 11947 | 11805 | 11787 | 453 | 484 | 439 | 489 | 5.01 |
| 06160 | ferredoxin | 95 | 118 | 73 | 67 | 3 | 4 | 5 | 4 | 4.94 |
| 00415 | NAD(P)-dependent malic enzyme | 484 | 533 | 516 | 514 | 26 | 25 | 24 | 26 | 4.79 |
| 00300 | DMT family transporter | 2498 | 2121 | 2259 | 1970 | 104 | 114 | 110 | 126 | 4.78 |
| 00275 | flavin monoamine oxidase family protein | 10295 | 13992 | 13600 | 13586 | 626 | 662 | 579 | 678 | 4.77 |
| 00325 | NADPH-dependent FMN reductase | 9914 | 9419 | 8818 | 9289 | 458 | 484 | 500 | 520 | 4.70 |
| 00285 | ester cyclase | 5059 | 4866 | 4905 | 4580 | 228 | 263 | 284 | 315 | 4.63 |
| 00315 | IS4 family transposase | 3661 | 4663 | 4203 | 5157 | 244 | 244 | 213 | 229 | 4.62 |
| 00305 | cupin domain-containing protein | 2790 | 2428 | 2551 | 2320 | 136 | 148 | 144 | 151 | 4.61 |
| 00270 | aldehyde dehydrogenase family protein | 8322 | 10768 | 10724 | 10364 | 534 | 583 | 539 | 613 | 4.59 |
| 06155 | NAD(P)/FAD-dependent oxidoreductase | 98 | 128 | 91 | 78 | 5 | 6 | 6 | 6 | 4.58 |

**Table S5 -** top 20 downregulated genes - Numbers in experimental columns represent transcripts per kilobase million (tpm) counts. Log2Fc = Log$_2$(Fold-change) between nicotine and glycerol growth conditions.

| Locus tag | Annotation | Nic-1 | Nic-2 | Nic-3 | Nic-4 | Gly-1 | Gly-2 | Gly-3 | Gly-4 | Log$_2$Fc |
|---|---|---|---|---|---|---|---|---|---|---|

| | | | | | | | | | | |
|---|---|---|---|---|---|---|---|---|---|---|
| 25185 | glycerol-3-phosphate dehydrogenase/oxidase | 1 | 1 | 2 | 2 | 643 | 665 | 651 | 643 | -8.27 |
| 25110 | YncE family protein | 20 | 19 | 16 | 20 | 3540 | 4482 | 3541 | 4233 | -7.31 |
| 05115 | extracellular solute-binding protein | 22 | 19 | 16 | 18 | 2766 | 3679 | 3659 | 3724 | -7.07 |
| 05120 | COG4315 family predicted lipoprotein | 35 | 28 | 27 | 32 | 3957 | 5207 | 5150 | 5537 | -6.93 |
| 15285 | glycerol-3-phosphate transporter | 6 | 7 | 6 | 7 | 1052 | 1033 | 980 | 1091 | -6.88 |
| 25195 | MIP/aquaporin family protein | 8 | 6 | 7 | 8 | 1057 | 1147 | 1034 | 1123 | -6.86 |
| 05125 | RNA polymerase sigma factor | 28 | 21 | 23 | 24 | 2157 | 2661 | 2594 | 2832 | -6.29 |
| 05130 | anti-sigma factor | 20 | 16 | 14 | 15 | 1345 | 1627 | 1575 | 1642 | -6.09 |
| 25190 | glycerol kinase GlpK | 18 | 17 | 17 | 18 | 1081 | 1159 | 1037 | 1092 | -5.54 |
| 14260 | YeiH family protein | 5 | 3 | 3 | 5 | 170 | 167 | 142 | 160 | -4.99 |
| 00875 | ATP-binding cassette domain-containing protein | 1 | 1 | 1 | 1 | 20 | 32 | 29 | 27 | -4.29 |
| 00870 | BC transporter permease subunit | 3 | 3 | 2 | 3 | 60 | 88 | 70 | 73 | -4.27 |
| 27980 | peptidoglycan DD-metalloendopeptidase family protein | 1 | 1 | 1 | 1 | 17 | 16 | 16 | 15 | -4.18 |
| 25115 | aminotransferase-like domain-containing protein | 1 | 3 | 1 | 0 | 27 | 33 | 36 | 36 | -4.04 |
| 03590 | 3D domain-containing protein | 46 | 33 | 32 | 32 | 943 | 764 | 703 | 765 | -4.00 |
| 00865 | FMNH2-dependent alkanesulfonate monooxygenase | 3 | 3 | 2 | 3 | 48 | 66 | 59 | 58 | -3.97 |
| 08845 | aspartate carbamoyltransferase catalytic subunit | 2 | 6 | 8 | 9 | 92 | 96 | 81 | 103 | -3.71 |
| 14240 | C40 family peptidase | 8 | 6 | 5 | 8 | 109 | 92 | 90 | 93 | -3.57 |
| 00860 | sulfonate ABC transporter substrate-binding protein | 6 | 5 | 6 | 5 | 71 | 106 | 88 | 90 | -3.52 |
| 18095 | competence type IV pilus minor pilin ComGG | 2 | 2 | 1 | 3 | 23 | 31 | 25 | 30 | -3.51 |

**Table S6**- RNA sequencing statistics. # of reads, and quality score statistics are from sequencing; reads assigned values are from FeatureCounts output and correspond to paired-end fragments.

| Sample ID | # Reads | Mean Quality Score | % Bases >= 30 | Reads assigned | Reads unassigned unmapped | Reads unassigned no features |
|---|---|---|---|---|---|---|
| nic-1 | 31,764,046 | 39.29 | 96.26 | 29,587,643 | 1,421,326 | 755,077 |
| nic-2 | 28,589,307 | 39.14 | 95.45 | 25,952,869 | 1,876,381 | 760,057 |
| nic-3 | 28,972,011 | 39.21 | 95.79 | 26,352,813 | 1,865,018 | 754,180 |

| | | | | | | |
|---|---|---|---|---|---|---|
| nic-4 | 31,335,139 | 39.21 | 95.84 | 28,335,084 | 2,196,148 | 803,907 |
| gly-1 | 34,145,852 | 39.26 | 96.1 | 31,668,815 | 2,014,805 | 462,232 |
| gly-2 | 30,984,011 | 39.18 | 95.66 | 29,047,401 | 1,528,064 | 408,546 |
| gly-3 | 30,886,213 | 39.21 | 95.76 | 29,093,886 | 1,390,108 | 402,219 |
| gly-4 | 37,704,419 | 39.35 | 96.57 | 35,617,669 | 1,557,382 | 529,368 |
